# Investigating the genetic diversity of H5 avian influenza in the UK 2020-2022

**DOI:** 10.1101/2022.12.03.518823

**Authors:** Alexander MP Byrne, Joe James, Benjamin C Mollett, Stephanie M Meyer, Thomas Lewis, Magdalena Czepiel, Amanda H Seekings, Sahar Mahmood, Saumya S Thomas, Craig S Ross, Dominic JF Byrne, Michael J McMenamy, Valerie Bailie, Ken Lemon, Rowena DE Hansen, Marco Falchieri, Nicola S Lewis, Scott M Reid, Ian H Brown, Ashley C Banyard

## Abstract

Since 2020, the UK and Europe, have experienced annual epizootics of high pathogenicity avian influenza virus (HPAIV). The first during autumn/winter 2020/21 involved the detected with six H5Nx subtypes although H5N8 HPAIV dominated in the UK. Whilst genetic assessment of the H5N8 HPAIVs within the UK demonstrated relative homogeneity, there was a background of other genotypes circulating at a lower degree with different neuraminidase and internal genes. Following a small number of summer detections of H5N1 in wild birds over the summer of 2021, autumn/winter 2021/22 saw another European H5 HPAIV epizootic, that has dwarfed the prior epizootic. This second epizootic was dominated almost exclusively by H5N1 HPAIV, although six distinct genotypes were defined. We have used genetic analysis to evaluate the emergence of different genotypes and proposed reassortment events that have been observed. The existing data suggests that the H5N1 circulating in Europe during late 2020, continued to circulate in wild birds throughout 2021, with minimal adaptation, but has then gone on to reassort with AIVs in the wild bird population. We have undertaken an in-depth genetic assessment of H5 HPAIVs detected in the UK, over the last two winter seasons and demonstrate the utility of in-depth genetic analyses in defining the diversity of H5 HPAIVs circulating in avian species, the potential for zoonotic risk and whether incidents of lateral spread can be defined over independent incursion of infection from wild birds. Key supporting data for mitigation activities.

**Importance:** High pathogenicity avian influenza virus (HPAIV) outbreaks devastate avian species across all sectors having both economic and ecological impacts through mortalities in poultry and wild birds, respectively. These viruses can also represent a significant zoonotic risk. Since 2020, the UK has experienced two successive outbreaks of H5 HPAIV. Whilst H5N8 HPAIV was predominant during the 2020/21 outbreak, other H5 subtypes were also detected. The following year there was a shift in subtype dominance to H5N1 HPAIV, but multiple H5N1 genotypes were detected. Through thorough utilisation of whole-genome sequencing, it was possible to track and characterise the genetic evolution of these H5 HPAIVs in UK poultry and wild birds. This has enabled us to assess the risk posed by these viruses at the poultry:wild bird and the avian:human interface and to investigate potential lateral spread between infected premises, a key factor in understanding threat to the commercial sector.

## Introduction

Since 2020, high pathogenicity avian influenza virus (HPAIV) outbreaks have devastated the poultry sector globally and constitutes a significant challenge to food security. Avian influenza viruses (AIVs) are classified as either low-pathogenicity (LP) or high-pathogenicity (HP) [1, 2]. LPAIVs generally cause mild infections, whilst HPAIVs can cause high mortality in a wide range of avian species. AIV subtypes are defined based on their surface glycoproteins, haemagglutinin (HA; H1-H16) and neuraminidase (NA; N1-N9) [3], but HPAIVs appear restricted to the H5 and H7 subtypes. In many countries legislation is in place for statutory control of H5 and H7 AIVs as notifiable animal pathogens [2] under the direction of the competent veterinary authority [4-6]. Commonly, national measure for HPAIV prevention and control focus on stringent biosecurity and depopulation of affected flocks with compensation to control of outbreaks [7]. As such, surveillance and monitoring of wild bird, and poultry populations for clinical signs is critical to detect and rapidly control such outbreaks [2]. Wild bird populations can maintain both HPAIVs and LPAIVs, and seasonal migration is considered a key factor in intercontinental dissemination of these viruses. Mixing of bird species at different sites enables genetic reassortment following coinfection, resulting in the emergence of novel AIVs [8]. Where infection pressure is high in birds and/or the environment, there is an increased risk of spread to poultry, as well as increasing the interface with other species, including humans (through occupational exposure) and scavenging animals [9-13]. However, the basis behind species-to-species adaptation events remains undefined.

AIVs are enveloped, negative-sense, single-stranded RNA viruses with each virion containing eight genome segments that together can generate up to 18 different proteins [14] including: the polymerase complex (polymerase basic protein (PB) 2 (PB2), PB1, and polymerase acidic protein (PA)); the nucleoprotein (NP)); the viral glycoproteins (HA and NA); structural proteins (matrix 1 (M1) and matrix 2 (M2)), and non-structural proteins (NS1 and NS2). Critically, the polymerase complex lacks proof-reading ability and so polymerase errors can occur, leading to genetic drift and subsequent maintenance of errors through successive generations. Ultimately, polymerase error rate drives viral evolution with these viruses, having an estimated error rate of up to 2.5×10^−4^ substitutions per nucleotide [15-17]. A further critical factor in the genetic evolution of these viruses is genetic reassortment following coinfection of the same cell. This feature of influenza virus biology can lead to dramatic genetic shifts, that can result in the emergence of novel influenza viruses, some with altered characteristics.

During the 2020/21 autumn/winter season, the United Kingdom of Great Britain and Northern Ireland, as well as the British Crown Dependencies (hereafter referred to as UK) and Europe experienced a significant AIV epizootic [18] with five H5 HPAIV subtypes (H5N1, H5N3, H5N4, H5N5 and H5N8), and at least 19 distinct genotypes being observed [19]. However, this multi-subtype epidemiological scenario changed dramatically during 2021/22 with the emergence of a dominant H5N1 HPAIV subtype, with only a small number of infections due to other subtypes (H5N2 and H5N8) reported [20]. Despite the dominance of the H5N1 subtype, significant genetic diversity was observed within these viruses across Europe. The initial detections in Europe possessed a HA gene with high similarity to that observed in the H5N1 viruses detected during the 2020/21 epizootic and into summer 2021 [21, 22]. This H5N1 sub-lineage, termed the B1 sub-lineage [21], is ancestral to those viruses that were detected in North America since late 2021 [23, 24]. Latter detections of H5N1 during the 2021/22 epizootic identified a second HA sub-lineage, B2, which encompassed viruses detected across Europe, and demonstrated divergence brought about by the accumulation of amino acid substitutions [21]. The divergence observed within the HA gene, was accompanied by additional diversity in the other seven influenza virus gene segments, resulting in a total of 16 genotypes by November 2021 [23].

In this study we generated whole-genome sequence (WGS) data for 240 AIVs from wild birds and poultry between 2020 and 2022. We have analysed outbreak cluster data to assess possible differentiation between independent incursions and interrogate the potential for lateral spread between infected premises.

## Methods

### Whole-genome sequencing

The samples obtained from H5 AIV positive investigations, (oropharyngeal or cloacal swab fluids, or tissue homogenates; brain, lung and trachea, intestines or mixed viscera), or virus isolates derived from these samples were used to generate whole-genome sequences (WGS). Virus isolates were obtained from clinical samples using 9- to 11-day-old specified pathogen free embryonated fowls’ eggs [2]. Total RNA was manually extracted, without the addition of carrier RNA from either clinical samples or viral isolates [25].

The extracted RNA was converted to cDNA using the SuperScript IV First-Strand Synthesis System with random hexamers (ThermoFisher), and then to double-stranded cDNA using the NEBNext Ultra II Non-Directional RNA Second Strand Synthesis Module (New England Biolabs). The double-stranded cDNA was then purified and concentrated using Agencourt Ampure XP beads (Beckman Coulter) and incubated at room temperature for 5 minutes and eluted in 10µL of 1M Tris-HCl pH 7.5 (Sigma), before quantification using the QuantiFluor dsDNA System (Promega). For preparation of the sequencing library, 1ng of purified dsDNA was used as the template and the library generated using the NexteraXT kit (Illumina). Sequencing libraries were run on either a MiSeq or NextSeq 550 (Illumina) with 2x150 base paired-end reads.

Raw sequencing reads were assembled using custom scripts: either FluSeqID (https://github.com/ellisrichardj/FluSeqID.sh) with consensus sequence generated using genconsensus.py (https://github.com/AMPByrne/WGS/blob/master/genconsensus.py), or denovoAssembly (https://github.com/AMPByrne/WGS/blob/master/denovoAssembly_Public.sh (accessed 21 September 2022). Some of the sequences used in this study were used in prior studies [9, 21, 26], but all sequences produced as part of this study are available through the GISAID EpiFlu Database (https://www.gisaid.org/) (**Table S1**).

### Phylogenetic analysis

Given the diverse nature of the H5Nx subtypes and genotypes observed in Europe during 2020-2022, it was important that an appropriate phylogenetic reference dataset was assembled to maintain the resolution of any subsequent analysis. To do this, all global AIV sequences from 2014-2022 were obtained from the GISAID EpiFlu Database and combined with the UK sequence data described in this study. This combined dataset was then used to generate phylogenies for each influenza gene segment using Nextstrain [27], and any duplicate sequences, sequences of poor quality, or demonstrating no topological relatedness to the UK sequences of interest were manually removed. For a minority of sequences, there remained a lack of ancestral sequences within the dataset. In these cases, the relevant sequences were used to query the GISAID EpiFlu BLAST Database to find similar sequences, which were then used to supplement the dataset.

Gene sequences were then aligned using Mafft v7.487 [28] and manually trimmed to the open-reading frame using AliView [29]. Phylogenetic trees were then inferred using the maximum-likelihood approach in IQ-Tree v2.1.4 [30] with ModelFinder [31] to infer the appropriate phylogenetic model and 1000 ultrafast bootstraps [32]. Ancestral sequence reconstruction and inference of molecular-clock phylogenies were performed using TreeTime [33]. Phylogenetic trees were visuallised using R version 4.1.1, with libraries ggplot2, ggtree [34] and treeio [35]. Phylogenetic incongruence analysis was performed using the maximum-likelihood phylogenetic trees using backronymed adapTable lightweight tree import code (BALTIC) as desribed previously [36]. Graphs were generated using Plotly version 5.8.0 (Plotly Technologies Inc.).

### Evaluation of viral polymorphisms associated with altered AIV characteristics

Viral protein sequences were screened for the presence of genetic polymorphisms that have been previously demonstrated to be associated with altered viral virulence, host tropism and antiviral resistance [37, 38] using a custom script: https://github.com/dombyrne/Influenza-Mutation-Checker, and database which is available upon request.

### Cluster analysis

Assessment of the potential for lateral spread versus independent introduction was undertaken on geographically linked cases, termed clusters. For each cluster, all wild bird and poultry detections, that were geographically and temporally relevant, and for which WGS data had been obtained, were included. Given that the different H5Nx genotypes were distinct, only sequences that were of the predominant genotype within the cluster sequences were included. These sequences were then concatenated to generate a single full-genome sequence covering all genes for each detection. The concatenated sequences were combined with a concatenated version of the genotype reference sequence (**Table 1**) and aligned as described above.

**Table 1.** H5Nx Subtypes and Genotypes identified through WGS in the UK during 2020-2022.

| Subtype | Genotypes Identified | Representative Virus | Total Sequences | HA Cleavage Site Motif | Sector |
| --- | --- | --- | --- | --- | --- |
| 2020/21 Epizootic |  |  |  |  |  |
| H5N1 HPAIV | 1 | A/chicken/England/043315/2020 | 3 | PLREKRRKR/GLF | Poultry<br>Wild Birds |
| H5N2 LPAIV | 1 | A/environment/England/030642/2020 | 1 | PQRETR/GLF | Poultry |
| H5N3 LPAIV | 1 | A/turkey/England/018179/2021 | 1 | PQRETR/GLF | Poultry |
| H5N3 HPAIV | 1 | A/peregrine falcon/Northern Ireland/AI102021-2/2021 | 1 | PLREKRRKR/GLF | Wild Birds |
| H5N5 HPAIV | 2 | H5N5.1:<br>A/Brent goose/England/095684/2020 | 1 | PQREKRRKR/GLF | Wild Birds |
|  |  | H5N5.2:<br>A/ mute swan/Wales/048068/2020 | 2 | PLREKRRKR/GLF |  |
| H5N8 HPAIV | 1 | A/Greylag goose/England/032698/2020 | 32 | PLREKRRKR/GLF (n=30)<br>PQREKRRKR/GLF (n=2) | Poultry<br>Wild Birds |
| 2021/22 Epizootic |  |  |  |  |  |
| H5N1 HPAIV | 6 | AIV07-B1: A/chicken/England/053052/2021 | 25 | PLREKRRKR/GLF | Poultry<br>Wild Birds |
|  |  | AIV07-B2:<br>A/Greylag goose/England/054503/2021 | 86 | PLREKRRKR/GLF | Poultry<br>Wild Birds |
|  |  | AIV08: A/chicken/Wales/053969/2021 | 2 | PLREKRRKR/GLF | Poultry<br>Wild Birds |
|  |  | AIV09: A/chicken/Scotland/054477/2021 | 82 | PLREKRRKR/GLF (n=78)<br>PLKEKRRKR/GLF (n=1)<br>PLREKRKKR/GLF (n=3) | Poultry<br>Wild Birds |
|  |  | AIV20: A/turkey/England/016515/2022 | 1 | PLREKRRKR/GLF | Poultry |
|  |  | AIV55: A/chicken/England/069816/2021 | 1 | PLREKRRKR/GLF | Poultry |
| H5N8 HPAIV | 1 | A/mute swan/England/298902/2021 | 1 | PLREKRRKR/GLF | Wild Bird |

Time-scaled phylogenetic trees were then inferred from the aligned concatenated sequences using BEAST version 1.10.4 [39] with the BEAGLE library [40]. The SRD06 nucleotide substitution model with a four-category gamma distribution model of site-specific rate variation and separate partitions for codon positions 1 + 2 versus position 3 with HKY substitution models on each with an uncorrelated relaxed clock with log-normal distribution, and the coalescent constant population size tree prior. For each cluster dataset, two independent Markov Chain Monte Carlo (MCMC) chains were run and combined using the LogCombiner tool in the BEAST package. Each chain consisted of 200,000,000 steps and was sampled every 20,000 steps and the first 10% of samples discarded as burn-in. The MCMC settings were chosen to achieve a post-burn-in effective sample size of at least 200. Discrete transition events between cluster detections were reconstructed using a symmetric continuous-time Markov Chain model with an incorporated Bayesian stochastic search variable selection (BSSVS) to determine which transition rates sufficiently summarised connectivity between detections [41]. SpreaD3 was used to visualise the rates of transmission through a Bayes factor (BF) test [42]. The BF represents the ratio of two competing statistical models, represented by their marginal likelihood, and in this case was used to determine the likelihood for transmission between detection events, as opposed to independent introductions [43]. The support of the BF for the transmission was interpreted as described previously [44]. Within each cluster, transmission events with a supporting BF of less than 3, or with supporting BF less than between any of the cluster sequences and the reference sequence, whichever was higher, were omitted.

## Results

### Incursions from wild birds drove the rapid emergence of the dominant H5N8 HPAIV subtype during autumn/winter 2020/21

Detection of HPAIV in Europe in autumn 2020 signalled that HPAIV was re-emerging [45], with the virus first being detected in the UK in a Greylag goose (*Anser anser*) in Gloucestershire on the 30^th^ October 2020. Wild bird detections during that season were limited to a 14-week period from the 30^th^ October to early February 2021, totalling 311 detections in Great Britain [46] with a further nine in Northern Ireland. Whilst multiple H5Nx HPAIV subtypes were detected across Europe, H5N8 dominated wild bird detections in the UK with 96% (n=292/320) of detections, whilst 4% were H5N1 (n=13/320), 2% were H5N5 (n=6/320) and 0.3% were H5N3 (n=1/320). The NA of the remaining samples were untyped (H5Nx; 3%, n=8/320). Additionally, 26 HPAIV-infected poultry premises were detected in the UK beginning with H5N8 HPAIV in Cheshire on 2^nd^ November. Twenty-three further H5N8 HPAIV detections, and two H5N1 HPAIV detections were made up to 31^st^ March 2021. Two notifiable LPAIV infections (H5N2 and H5N3) were also detected on poultry premises.

For H5N8 HPAIV detections in wild birds (**Figure 1A**) and poultry (**Figure 1B**) in the UK, WGS data demonstrated that all sequences were highly identical (>98.1%) across all gene segments, suggesting a single H5N8 genotype. Phylogenetic analysis of the UK H5N8 HA (**Figure 2 and Figure S1A**), and other gene segments (**Figure S1B-H**), demonstrated high similarity to viruses detected in Europe during the same 2020/21 epizootic period, and implicated a single common H5N8 HPAIV ancestor, A/chicken/Iraq/1/2020, detected in May 2020. HPAIVs resulting from this ancestral strain were likely responsible for spread across Europe, as well as the Middle East and Central Asia. The HA cleavage site (CS) motif of the UK H5N8 sequences was PLREKRRKR/GLF, with only two sequences showing any differences (both P**Q**REKRRKR/GLF) (**Table 1**) [47].

**Figure 1.**
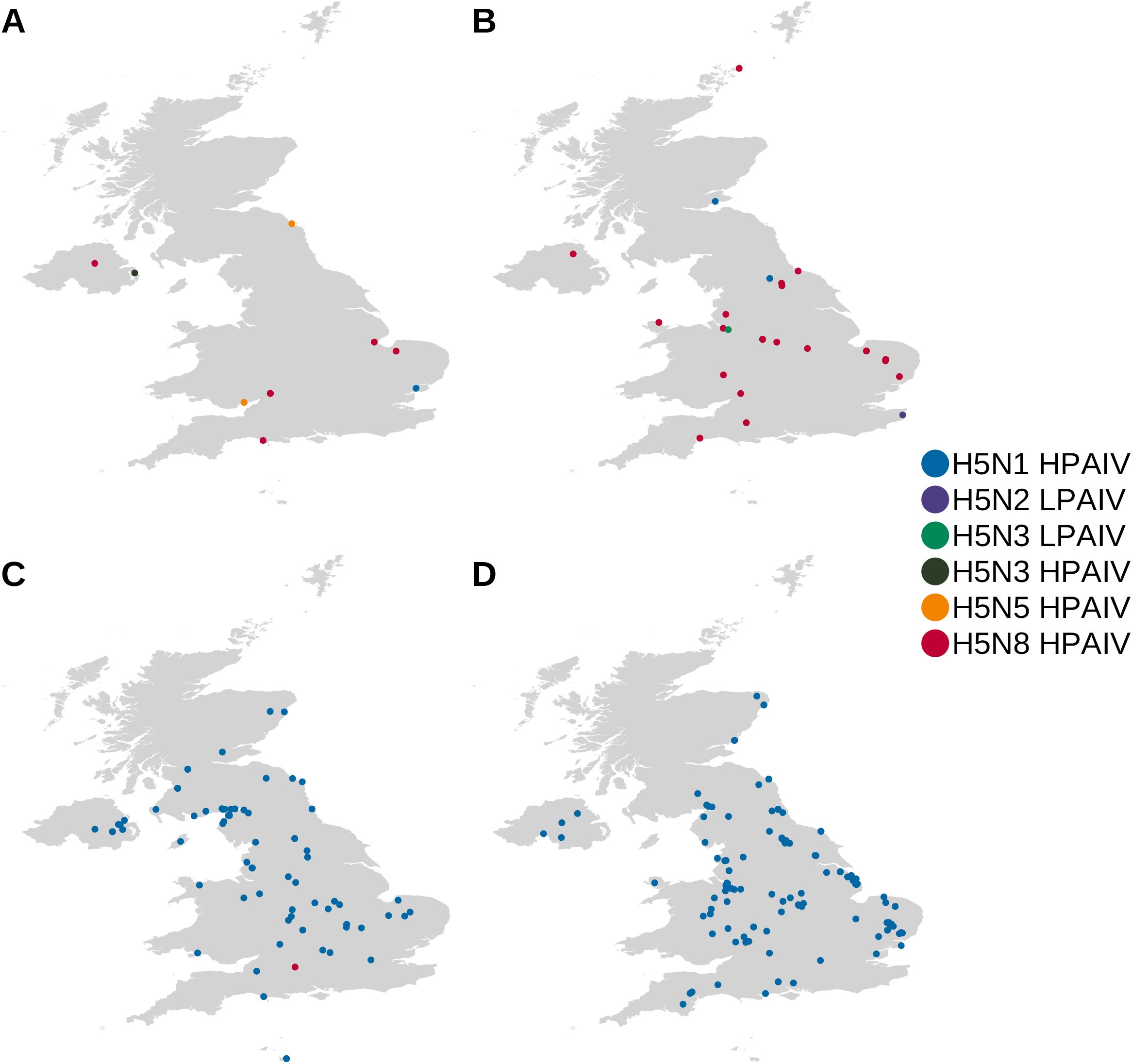
Geographic distribution of H5Nx AIVs that were sequenced during 2020-2022. Geographic distribution of H5Nx AIV viruses that were sequenced from wild birds (A and C) and poultry (B and D) during the 2020/21 (A and B) and 2021/22 (C and D) epizootics in the UK. Locations are coloured according to AIV subtype and pathotype.

**Figure 2.**
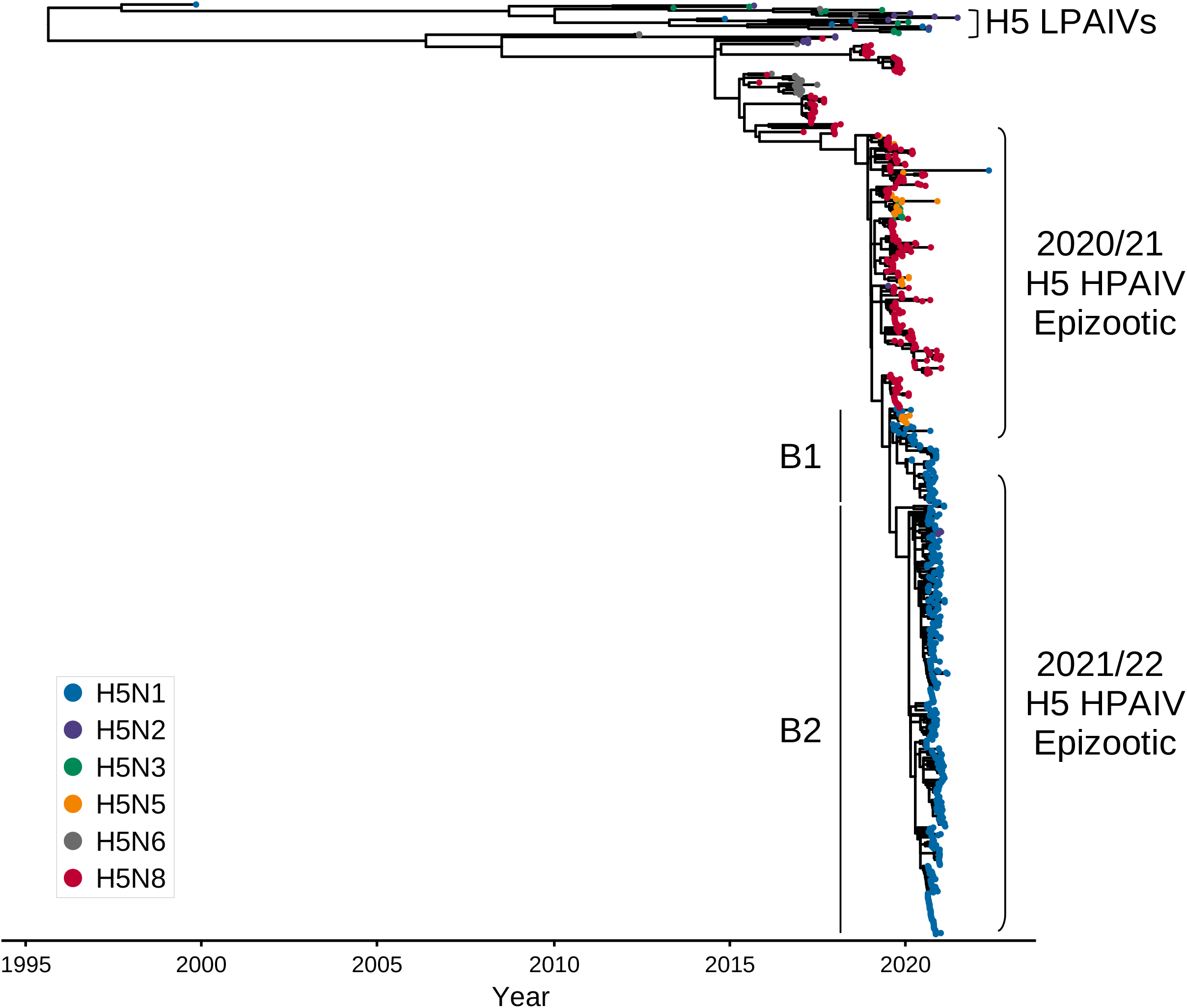
The HA of the H5Nx HPAIVs from 2020-2022 were derived from a common ancestor. Time-resolved maximum-likelihood phylogenetic tree of the HA gene from H5Nx AIVs collected from the UK between 2020-2022, with relevant global reference sequences. The tips are coloured according to viral subtype and the sequences obtained from either the 2020/21 and 2021/22 H5 HPAIV epizootics are indicated. For the H5N1 HPAIV sequences the B1 and B2 sub-lineages are also shown.

H5N5 HPAIV was only detected in wild birds in the UK, and three genome sequences were generated from samples collected (**Table 1**). However, even within this small number of sequences, two distinct H5N5 genotypes were identified through phylogenetic analysis. Both genotypes derived the majority of their gene segments from A/chicken/Iraq/1/2020, similar to the H5N8 HPAIV genotype observed in the UK (**Figure S1A-H**) and shared the same two HA CS motifs (**Table 1**). The N5 gene appeared to be obtained through reassortment with local AIVs as it demonstrated similarity with H5N5 AIVs detected in Europe in 2020 (**Figure S1B**). However, the two UK H5N5 HPAIV genotypes differed in the PA gene. Whilst H5N5.1, represented by A/Brent goose/England/095684/2020, detected in Northumberland (**Figure S2A**), had a PA gene that was highly similar to that of A/chicken/Iraq/1/2020, H5N5.2, represented by A/mute swan/Wales/048068/2020, had a different PA segment closely related to those of AIVs detected in Eurasia, indicating potential reassortment (**Figures 3A, 3B and S1E**). Nevertheless, this PA segment was also observed in H5N5 AIVs identified in European wild birds and poultry in 2020/21.

**Figure 3.**
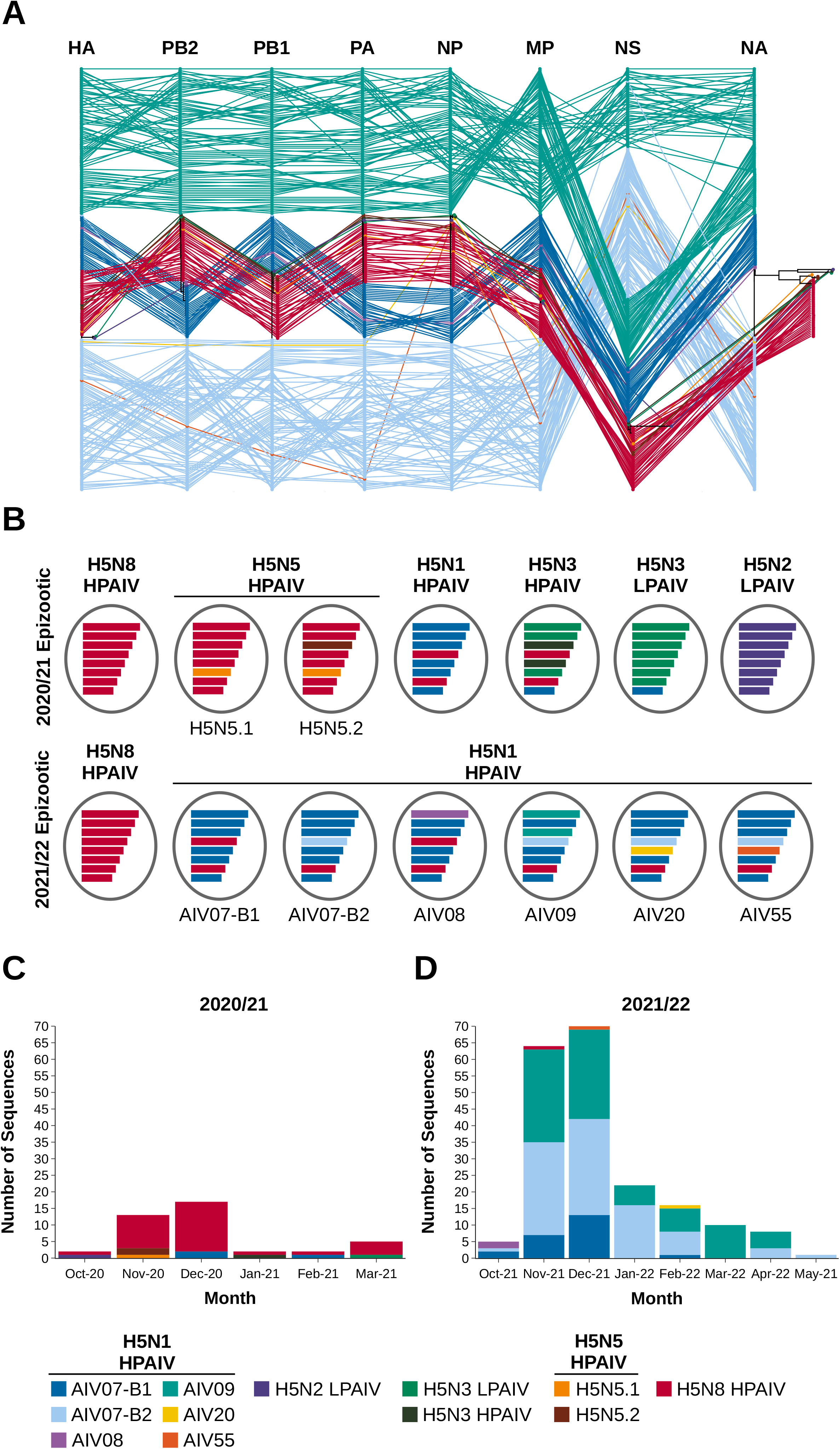
H5Nx AIVs from the UK collected between 2020-2022 demonstrate wide genotypic diversity. (A) Phylogenetic incongruence analysis of H5Nx sequences from the UK from AIVs collected between 2020-2022. Maximum-likelihood phylogenetic trees for all gene segments from equivalent strains are connected across the trees, with tips and connecting lines coloured according to genotype. (B) Schematic representation of the different H5Nx genotypes from the UK between 2020-2022. It should be noted that whilst the HA gene of the H5N1 HPAIV B2 sub-lineage, is coloured differently for the purposes of this diagram, it is still derived evolutionarily from the A/chicken/Iraq/1/2020 H5N8 HPAIV HA gene. (C and D) Number of sequences for each UK H5Nx genotype generated during the 2020/21 and 2021/22 epizootics, respectively.

The third HPAIV subtype detected during the 2020/21 epizootic, H5N1, was only identified in England and Scotland. These H5N1 HPAIVs, like those also observed in Europe, demonstrated similarity in the HA (**Figures 2 and S1A**) and matrix protein (MP) gene segments with A/chicken/Iraq/1/2020 H5N8 HPAIV (**Figure S1G**). However, the other gene segments were highly identical to H5N1 sequences detected throughout Europe, and Africa from 2020-2021, with relatedness to Eurasian AIV sequences as far back as 2016, resulting in a singular genotype (**Figures 3A and 3B**).

H5N3 HPAIV was detected in the UK in a single peregrine falcon (*Falco peregrinus*) from Northern Ireland (**Table 1**) being characterised as a reassortant including the HA (**Figure 2 and S1A**) and MP (**Figure S1G**) from A/chicken/Iraq/1/2020 (**Figure 3A and 3B**) and remaining genes from Eurasian LPAIVs. Interestingly the H5N3 HPAIV NS gene segment had greater than 97% identity to the H5N1 HPAIV sequences.

The two LPAIVs were detected in England during 2020/21 and included an H5N2 virus isolated from faecal material in Kent from a mixed poultry premises and an H5N3 from turkeys in Cheshire. The H5N2 LPAIV was genetically similar to other H5N2 sequences obtained from Europe and Asia during the same period (2020-2021) (**Figure S1A-H**). The H5N3 LPAIV showed limited similarity to the other sequences obtained from the UK during this period although the PB2, PB1 and NS segments clustered with those of the UK H5N3 HPAIV sequence.

### Re-emergence and dominance of H5N1 HPAIV during 2021/22 season

Re-emergence of H5N1 HPAIV started following detection within the Great skua (*Stercorarius skua*) population on the Shetland Islands off the north coast of Scotland during summer 2021 [22] and was detected again in the UK in wild birds and poultry from October 2021. The first poultry case occurred in Worcestershire on 26^th^ October 2021, with the first wild bird detection made in a gull (*Larus canus*) collected on 14^th^ October 2021 from Scotland through the UK passive surveillance system. From these initial incursions, until May 2022, over 1,000 wild birds tested positive for H5N1 HPAIV across the UK, with significant impact on the UK poultry sector involving infection of over 115 premises. All poultry cases involved infection with the H5N1 virus that had circulated at a lower frequency during 2020/21.

During the 2021/22 epizootic, WGS of 196 viruses obtained from poultry and wild birds (**Figures 1C and 1D**) in the UK demonstrated the presence of six distinct genotypes (**Table 1**). These genotypes were based on identity to the progenitor H5N1 HPAIV detected in the previous year (2020/21) and found in the Great skua population; genotypes are denoted based on the first detection in wild birds or poultry (**Figures 3A and 3B**).

The first H5N1 genotype detected was AIV07, which had the same gene constellation as the H5N1 detected across wild birds and the two UK poultry cases during the previous epizootic (**Figures 3A, 3B and S1A-H**). However, this genotype initially demonstrated divergence within the HA gene [21] (**Figure 2**) and has been defined as two separate genotypes. The AIV07-B1 genotype contained a HA with high similarity to the virus from 2020/21 and was the primary UK H5N1 detection during 2021/22. However, AIV07-B1 later became a minority population in the UK and was not detected after February 2022 (**Figure 3C**). The AIV07-B2 genotype, possessed a HA gene that had diverged from AIV07-B1 [21] (**Figure 3C**), although both genotypes were detected in wild bird and poultry cases throughout the UK (**Figure S2C and S2D**).

The third H5N1 genotype, AIV08, was only detected in a single poultry case, and an associated wild bird detection from the same site in Wales in October 2021 (**Figure S2C and S2D**). The AIV08 genotype, shared high genetic similarity in all gene segments to AIV07-B1, except for the PB2 segment (**Figures 3A and 3B**), which had high similarity to that observed in LPAIVs detected in the Netherlands and the Republic of Ireland since 2020 (**Figure S1C**). H5N1 HPAIV sequences with a similar PB2 segment were detected in poultry and wild birds in France, Italy, Moldova and Romania between October 2021 and February 2022.

The fourth H5N1 genotype, AIV09, was the second most prevalent (**Table 1 and Figure 3C**), and was first detected in Scotland in November 2021, but has since been detected across the UK in poultry and wild birds (**Figure S2C and S2D**). The AIV09 genotype shared the PB1, NP, NA, MP, and NS segments with AIV07-B1 and AIV07-B2 but possessed the HA from the B2 sub-lineage. The PB2 and PA segments, however, demonstrated dissimilarity to both AIV07 genotypes, as well as AIV08 (**Figures 3A and 3B**). The AIV09 PB2 showed high genetic similarity to that seen in the H5N3 AIVs in the UK and Europe during the 2020/21 epizootic (**Figure S1C**), and the PA with LPAIVs from the Netherlands and Belgium detected since 2017 (**Figure S1E**). Interestingly, phylogenetic incongruence analysis suggests that there may be two separate lineages within the AIV09 genotype, based on differences in the NS segment (**Figure 3A**). However, the nucleotide identity of all the UK H5N1 sequences from 2020-2022 share greater than 98.21% identity for the NS gene and the topography of the phylogenetic tree demonstrates that the European H5N1 NS genes were derived from a single common ancestor (**Figure S1H**). Therefore, it can be inferred that the NS gene is the same across the AIV09 genotype sequences.

The final two H5N1 HPAIV genotypes, AIV20 and AIV55, were only detected once during the study period (October 2020 to May 2022). AIV20 was detected on a turkey farm in Lincolnshire in February 2022, whilst AIV55 was detected in chickens from County Durham in December 2021 (**Figures 3C and S2D**). Both genotypes shared seven of their eight gene segments with AIV07-B2, including the HA gene, but had alternative NP gene segments (**Figure 3A and 3B**). For AIV20, the NP segment demonstrated similarity to those from AIVs in the Netherlands and Belgium, but also the H5N3 HPAIVs observed during 2020/21 (**Figure S1F**). The AIV55 NPsegment was more closely related to those observed in H5N1 and H5N5 HPAIVs from Eastern Europe and Russia, as well as a H12N5 sequence from Belgium.

Finally, whilst no detections of H5N8 HPAIV were made in poultry during the 2021/22 season, this subtype was found in a single Mute Swan (*Cygnus olor*) collected from Wiltshire in November 2021 (**Figure 1C**). The virus sequence demonstrated high similarity to the H5N8 observed in the UK during the 2020/21 epizootic and was the same genotype (**Figures 3A, 3B and S1A-H**).

### Evaluation of host tropism markers in H5Nx sequences

In accordance with standard risk assessments within the UK, AIV sequences obtained from outbreaks were assessed for the presence of previously defined zoonotic molecular markers associated with increased virulence, alterations in host tropism and resistance to antivirals [37, 38] (**Table S2**). A numbering of polymorphisms were identified, including the HA T156A substitution, which is associated with increased binding to α2-6-linked sialic acids [38, 48, 49]. Interestingly, the PB1 D3V substitution, was identified within the majority of sequences and genotypes assessed but was differentially identified between the two H5N5 genotypes; the substitution was present in the H5N5.1 genotype, whilst it was absent from H5N5.2. This may indicate genetic drift between the two genotypes but cannot be confirmed given the limited number of H5N5 HPAIV sequences obtained. The H5N5 and H5N8 sequences were also exclusively found to possess a truncated PB1-F2 protein, consisting of only 11 amino acids, whilst all other sequences had a full-length (90 amino acid) protein.

The M2 A30S amino acid substitution, associated with reduced susceptibility to amantadine and rimantadine [38, 50-54] was identified in a single H5N1 sequence, whilst the NA I117T substitution shown to reduce susceptibility to NA-inhibitors [38, 55] was identified in all H5N2 and H5N3 sequences. The PA I38T substitution associated with reduced baloxavir susceptibility [56] was not identified in any of the sequences analysed.

### Genetic assessment of outbreak clusters to assess the likelihood of lateral spread

During the 2020/21 epizootic, the H5N8 sequences characterised shared high genetic identity and formed a single genotype. The geographical distribution of cases in the UK during 2020/21 suggested that there was no direct epidemiological relationship between infected premises (IPs) and supports the likelihood of multiple independent primary introductions from wild birds in each instance. Movement of birds prior to the development of disease may, of course, have facilitated outbreaks in geographically distinct areas, but evidence to evaluate this could not be drawn from genetic data. One exception to this was two closely linked IPs located in North Yorkshire (Cluster 1), which were confirmed to be H5N8 HPAIV positive within four days of each other and epidemiologically defined pathways demonstrated (data not shown) (**Table S3**). A Bayesian stochastic search variable selection (BSSVS) analysis was used to identify well-supported rates of transition between IPs and support was quantified using Bayes Factors (BF) (**Figure 4 and Table S4**). This analysis demonstrated that there was no strong BF support for transmission between premises, suggesting separate introductions onto both premises.

**Figure 4.**
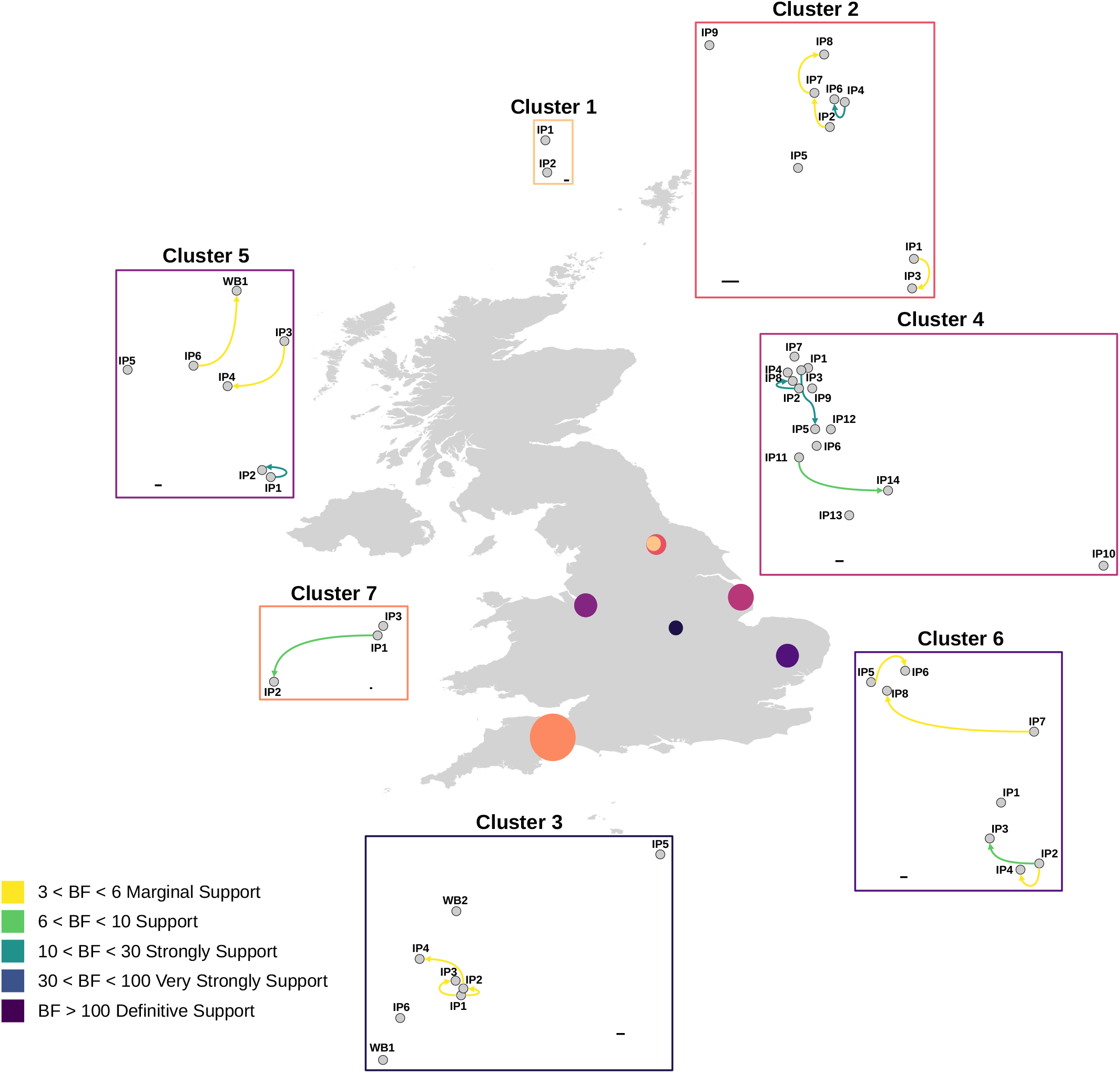
Analysis of H5Nx sequences suggests limited lateral transmission between geographically related HPAIV detections. Outputs of the BSSVS analysis for the seven geographical clusters of H5Nx HPAIV detections investigated for the potential of lateral transmission to have occurred. Each geographical cluster is represented by a separate network diagram using the relative location of each infected premises (IP) or wild bird (WB) detection. Arrows are coloured according to the relative strength, inferred using a Bayes Factor (BF), by which the transmission rates are supported. Scale bars are provided for each cluster, representing 1 km.

The escalation in cases during the 2021/22 epizootic led to further investigations into the potential for lateral spread between IPs. Six geographically linked groups of premises, or ‘clusters’, were detected within short timeframes of each other in poultry dense regions of England [57]. The presence of multiple H5N1 HPAIV genotypes in circulation within the UK enabled distinction of independent incursion wherever different genotypes were detected. This also enabled refinement of the IPs that constituted each cluster based on the major genotype detected. A BSSVS analysis approach was then applied to this refined set of IPs to assess the potential for lateral spread as opposed to independent introductions. Of the six clusters investigated from the 2021/22 epizootic, all but one involved the AIV09 genotype, with only Cluster 5 involving the AIV07-B2 genotype (**Table S3**).

Cluster 2 consisted of nine IPs, where AIV was detected between 12^th^ November and 8^th^ December 2021, including a mixture of chicken and turkey premises. The BSSVS analysis suggested potential for lateral spread between IP4 and IP6 (both turkey premises) with strong BF support (**Figure 4 and Table S5**).

Cluster 3 involved six IPs (five chicken and one turkey premises), and two wild bird detections; a mute swan and a common gull, that were collected between 13^th^ November 2021 and 8^th^ January 2022 in Leicestershire. Interestingly, using this approach neither wild bird sequence was proposed to be the origin of H5N1 for the poultry premises within this cluster. BSSVS analysis suggested that IP1 was a potential source of virus for both IP2 and IP3, with the former being linked to IP4 also. Furthermore, these IPs were all chicken premises located in close proximity. The BF support for lateral transmission was low for all other IPs in this cluster, suggestive of independent introductions from wild birds directly or indirectly (**Figure 4 and Table S6**).

Cluster 4 was the largest geographic cluster investigated, involving 14 IPs; chickens (n=10), ducks (n=1), turkeys (n=2) and one IP housing chickens and ducks (IP12), that were detected between 11^th^ December 2021 and 8^th^ January 2022 in Lincolnshire. However, there appeared to be only two strongly supported transmission events following BSSVS analysis: between IP3 and IP5, and between IP2 and IP8, all four of which were chicken premises. Interestingly, whilst IP2 and IP8 were located close together, but IP3 and IP5 were more distant, with several other premises situated between them in the line-of-flight. The BSSVS analysis suggested that the remaining IPs were likely the result of independent introductions (**Figure 4 and Table S7**).

Cluster 5 involved six poultry IPs (one chicken, one duck and four turkey premises) and a wild bird detection (a mute swan) confirmed between 17^th^ December 2021 and 27^th^ January 2022 in Cheshire. Within this cluster, the transmission from IP3 to IP4 (both turkey premises) had marginal support, as did transmission from IP6 to WB1 (**Figure 4 and Table S8**). However, given that WB1 was detected over a month before IP6 (**Table S3**), this may indicate that both detections were the result of a singular, unidentified introduction source, most likely another wild bird. Only the transmission between IP1 and IP2, which were in close proximity was strongly supported by the BSSVS analysis.

Cluster 6 consisted of eight IPs (one goose, five duck and two chicken premises) confirmed between 25^th^ February and 4^th^ April 2022 in Suffolk. The BSSVS analysis suggested transmission from IP2 to IP3, and IP4, as well as IP5 to IP6, and IP7 to IP8 (**Figure 4 and Table S9**). However, these transmissions were supported by low BFs, with the infection of remaining premises proposed to be independent introductions.

Cluster 7 was the last cluster to be identified and investigated; consisting of three premises (one containing both ducks and geese, along with one duck and one chicken premises) detected between the 4^th^ and 12^th^ April 2022 in Devon. The BSSVS analysis suggested that IP1 may have transmitted virus to IP2, which were the most distant IPs geographically, however, the support for this was low (**Figure 4 and Table S10**).

## Discussion

The European HPAIV epizootic during 2020/21 resulted in a total of 3,555 virus detections across 28 European countries [18]. Whilst H5N8 predominated during that epizootic (88% of total detections), multiple other subtypes were detected including: H5N5 in captive birds, poultry, and wild birds (3% of total detections); H5N1 in poultry and wild birds (3% of total detections); and H5N3 and H5N4 in wild birds (1% and 0.5% of total detections, respectively) with the majority being detected between 5^th^ October 2020 and 23^rd^ February 2021 [18]. In the UK, there was a total of 26 positive poultry premises and 320 wild bird positives, with the majority of cases being H5N8, followed by H5N1, H5N5 and H5N3. H5N4 HPAIV, whilst detected in Germany, the Netherlands and Switzerland, was not detected in the UK [18]. The high degree of genetic relatedness to A/chicken/Iraq/1/2020 across H5 HPAIV subtypes supports the hypothesis that H5N8 was introduced into Europe via a single common progenitor, most likely during late 2020 via Russia and Eastern Europe [58], although migratory movements driving emergence remain undefined. Regardless of the mechanism of introduction, multiple H5 HPAIV subtypes were detected with significant genotypic diversity [59]. Critically, whilst reassortants involving some internal genes have been described, the HA and MP gene segments were conserved throughout the European H5 HPAIV detections in 2020-2021 [59]. Furthermore, in contrast with genetic diversity observed in Europe [18], in the UK, genotypic diversity was limited to single H5N8, and two H5N5 genotypes during the 2020/21 epizootic. The observed homogeneity in UK genotypes is hard to explain but may be a factor of partial immunity in ducks preventing coinfection that might lead to reassortment, alongside rapid lockdown of premises testing positive to minimise the risk of lateral spread.

During 2020/21, two H5 LPAIVs (H5N2 and H5N3) were detected in unrelated poultry cases in the UK. From a genetic standpoint, the H5N3 LPAIV contained three gene segments with high sequence identity to the H5N3 HPAIV detected in Northern Ireland in January 2021. Detection of notifiable LPAIVs often relies upon serological flock assessment and rapid statutory follow-up investigation of premises where H5 or H7 specific antibodies are detected. These detections were both the result of this testing algorithm, and neither of the LPAIVs detected in these instances caused any overt clinical disease in the birds involved. A paucity of data underscores our lack of understanding with respect to LPAIV circulation. However, having the environment for the interaction between species that might transmit both LPAIVs and HPAIVs is critical to coinfection events and factors, including susceptibility and prior immunity, that drive this.

The last detection of H5N1 HPAIV in poultry in the UK was in late March 2021, with the last wild bird detection in April. In contrast, detection of HPAIV across northern and eastern Europe in poultry, wild and captive birds continued through to May 2021 [18]. For the first time, summer detections of H5N1 HPAIV occurred following emergence in Great skuas off the north coast of Scotland during July and August 2021 [22] with the virus being closely related to the H5N1 HPAIV detected in the UK and Europe during the 2020/21 epizootic. This virus was also detected a further 54 times during summer 2021 in wild birds from Europe (Estonia, Germany, Finland, Latvia, the Netherlands and Sweden) [19], suggesting maintenance of this virus across wild bird populations [22].

Within the UK, the detection of two sub-lineages of the AIV07 genotype: AIV07-B1 and AIV07-B2 occurred between October 2021 and May 2022. The AIV07-B1 genotype was detected in 12 poultry IPs and 10 wild bird detections, whilst AIV07-B2 was detected in 42 poultry cases and 44 positive wild birds during the same period. The AIV07-B1 genotype was also detected in the human case of H5N1 HPAIV infection during December 2021 although no evidence of mammalian adaptation was observed [26]. Whilst the H5N1 B1 sub-lineage has been detected across North America since the end of the 2020/21 European HPAIV epizootic [23, 24] it has been a minor sub-lineage detected in Europe during the 2021/22 epizootic (UK (n=25), France (n=1), Germany (n=7), Republic of Ireland (n=5), Sweden (n=7) and Denmark (n=1). Re-emergence of HPAIV in the UK is hypothesised to have occurred via two routes: i) the AIV07-B1 genotype was likely introduced from Sweden and Denmark whilst; ii) the AIV07-B2 genotype was likely introduced from Northern Europe having likely originated in Russia and Eastern Europe. In both cases, the virus was likely introduced following the movements of migratory waterfowl, although local asymptomatic circulation in local wild bird populations cannot be excluded.

The detection of the AIV08 genotype in only a single poultry case and a single associated wild bird case during the 2021/22 season is of interest. This genotype had undergone reassortment of the PB2 segment, with that segment being most closely related to that of European AIVs of varying subtypes detected in poultry and wild birds since 2018. The clustering of AIV08 HA with the H5N1 B1 sub-lineage suggests that it may have emerged following a reassortment event between an AIV07-B1 virus and an undefined AIV present within the wild bird population. Its apparent extinction in the UK and limited detection of AIVs containing a similar PB2 segment to this genotype across Europe may indicate poor segment compatibility, perhaps resulting in reduced viral fitness or different host tropism.

The third genotype detected within the UK, AIV09, has high sequence identity with the AIV07 genotypes but contains different PB2 and PA genes. The PB2 segment of AIV09 had high sequence similarity with European H5N3 sequences (LPAIV and HPAIV) detected during 2020-2021, as well as LPAIV subtypes detected in Eurasian poultry and wild birds. Similarly, the PA gene has high similarity with those described in LPAIVs detected in Belgium and the Netherlands from 2017 to 2019, and more distantly with H5N5, H5N3 HPAIVs and H5N2 LPAIV from the 2020/21 epizootic. Reassortment facilitated through interactions between wild birds at, or on route to their breeding grounds during summer 2021 likely also enabled the emergence of this genotype. The AIV09 genotype is presumed to have been introduced into the UK from the east, due to the relatedness to contemporary H5N1 viruses detected in late summer in Russia.

The AIV20 and AIV55 genotypes, shared substantial similarity to the AIV07-B2 genotype, except for their NP genes, which were closely related to wild bird AIVs detected in Belgium in 2017 and 2020, respectively, and are distinct from the NP observed in the other UK genotypes detected during 2020-2022. These genotypes may have followed a similar migration pathway to the AIV07-B2 genotype, but potentially obtained their novel NP genes through reassortment with AIVs circulating in European wild birds. Critically, a paucity of viral sequence data, particularly for LPAIVs, means that conclusions around the exact emergence pathways for these viruses remain unclear.

Assessment of the sequences generated in this study for polymorphisms associated with increased virulence, altered host tropism or antiviral resistance found there was no association between H5 genotype and the observed polymorphisms. Previous studies have demonstrated that adaptive changes occur within the polymerase complex following mammalian infection but that the change identified (PB2 D701N) was most likely a single mutation that, alongside other mammalian adaptations may increase zoonotic threat [9]. Similarly, the PB2 E627K shown to be involved in adaptation to mammalian hosts [38, 60-64], and considered a significant marker of mammalian adaptation, was only identified in a single H5N1 sequence obtained from poultry. Nevertheless, the risk of infection at the poultry-human interface posed by these H5Nx clade 2.3.4.4b viruses remains low, as evidenced by the low number of human infections that have been detected globally since 2020 [10], despite the substantial infection pressure and potential for opportunistic infections at the avian-human interface during the concurrent epizootics.

The apparent maintenance of H5N1 within wild bird species during the summer months of 2021 in Northern Europe is a key shift in epidemiology compared to what has been previously observed with clade 2.3.4.4b H5 HPAIVs. Certainly, in Europe this is the first time that H5 HPAIV maintenance has been observed in wild birds and this likely facilitated genetic diversification, through local coinfection and reassortment with AIVs enzootic in the wild bird population. The apparent stability of the different H5N1 genotypes, following introduction into the UK in late 2021, is clearly demonstrated by genotype distribution across overlapping locations and disparate species. As before, defining the origin of these genotypes is problematic without a greater understanding of the circulation of HPAIV in species that tolerate infection in the absence of disease, and LPAIVs amongst all bird species.

The genetic analysis of different viruses from geographically linked clusters aimed to define where independent incursion may have occurred over the likelihood of lateral spread due to inefficient biosecurity practices. The determination of multiple H5N1 genotypes during the 2021/22 outbreak enabled, at least where different genotypes were observed, some conclusions to be made at the consensus level, although genetic divergence could not conclusively be used to differentiate between introduction sources, and a more rigorous approach was required [65-68]. Investigations of this type will become more important in understanding incursion risks and factors driving virus spread. Certainly, the utility of WGS in characterising outbreaks is critical and comprehensive genetic data, particularly for LPAIVs, with a deeper level of analysis would benefit assessments of this type.

In conclusion, the change in HPAIV epidemiology and maintenance within local populations raises uncertainties in defining risk of incursions. Interestingly, the two UK epizootic events appear to have demonstrated differential plasticity in HA/NA interactions with the 2020/21 H5 successfully interacting with multiple NA types, whilst the 2021/22 H5 has exhibited an apparent preferential interaction with N1 that has facilitated proliferation across a broader range of species than seen previously. Furthermore, the replication fitness of these viruses appears to have a tolerance for reassortment of several segments, particularly the polymerase complex. A rapid evaluation of factors influencing the impact of genotype on phenotype is required to better understand virus host interactions.

## Data availability

All sequence data generated and used in this study are freely available through GISAID EpiFlu Database (https://www.gisaid.org/). All accession numbers are provided in **Table S1**.

## Acknowledgements and Funding

The authors would like to thank the National and International Reference Laboratory staff, as well as the Central Unit for Sequencing and PCR at APHA for their assistance with this study. The authors would also like to thank Nadja Howton-Cheney (APHA) for their help with refining the database used for assessing viral changes associated with altered virulence. Finally, the authors would like to thank Samantha Lycett (University of Edinburgh) for their advice and guidance with respect to the Bayesian analyses.

We acknowledge the authors, originating and submitting laboratories of the sequences from GISAID’s EpiFlu Database on which this research is based, and analyses described in text. All submitters of the data may be contacted directly via the GISAID website (www.gisaid.org).

This work was funded by the Department for Environment, Food and Rural Affairs (Defra, UK) and the Devolved Administrations of Scotland and Wales, through the following programmes of work: SV3400, SV3032, SV3006 and SE2213. Funding for diagnostic testing in Northern Ireland was provided by the Department for Agriculture, Environment and Rural Affairs (DAERA). The writing and data analysis for this manuscript was also supported, in part by the “DELTA-FLU” project funded by the European Union’s Horizon 2020 research and innovation program under grant agreement no. 727922. ACB, JJ and IHB were also part funded by the BBSRC/Defra funded research initiative ‘FluMAP’ (BB/X006204/1).

## Ethical statement

All samples were obtained from dead animals collected as part of the epizootic.

## Conflicts of Interest

The authors declare no conflicts of interest.

## Figure Legends

**Figure S1**. Time-resolved maximum-likelihood phylogenetic trees containing the H5Nx sequences obtained from the UK, with relevant global reference sequences. (A) H5, (B) NA, (C) PB2, (D) PB1, (E) PA, (F) NP, (G) MP and (H) NS. Sequences are coloured according the H5 subtype, and UK H5Nx genotypes are illustrated. The sequences obtained from the UK are indicated with circular tip shapes.

**Figure S2**. Geographic distribution of H5Nx AIVs that were sequenced from wild birds (A and C) and poultry (B and D) during the 2020/21 (A and B) and 2021/22 (C and D) epizootics in the UK. Locations are coloured according to AIV subtype, pathotype and genotype.

**Table S1**. GISAID EpiFlu accession numbers for all sequences generated in this study.

**Table S2**. All polymorphisms associated with altered virulence, host susceptibility and antiviral resistance for the different H5 AIV subtypes detected in the UK between 2020-2022. All different polymorphisms observed for each subtype are shown, along with their previously demonstrated phenotype.

**Table S3**. Sequences used to investigate lateral spread between infected premises (IP) and wild birds (WB) in the different geographic clusters. The collection date and associated H5 subtype/genotype, as well as the reference sequence used for each cluster are also provided.

**Table S4**. Outputs of the BSSVS analysis for Cluster 1 showing the rates of transmission between sampled infected premises (IP).

**Table S5**. Outputs of the BSSVS analysis for Cluster 2 showing the rates of transmission between sampled infected premises (IP).

**Table S6**. Outputs of the BSSVS analysis for Cluster 3 showing the rates of transmission between sampled infected premises (IP) and/or wild birds (WB).

**Table S7**. Outputs of the BSSVS analysis for Cluster 4 showing the rates of transmission between sampled infected premises (IP).

**Table S8**. Outputs of the BSSVS analysis for Cluster 5 showing the rates of transmission between sampled infected premises (IP) and/or wild birds (WB).

**Table S9**. Outputs of the BSSVS analysis for Cluster 6 showing the rates of transmission between sampled infected premises (IP).

**Table S10**. Outputs of the BSSVS analysis for Cluster 7 showing the rates of transmission between sampled infected premises (IP).

## Notes

### Competing Interest Statement

The authors have declared no competing interest.

## References

1. Alexander, D.J. and I.H. Brown, History of highly pathogenic avian influenza. Revue scientifique et technique (International Office of Epizootics), 2009. 28(1): p. 19–38.

2. OIE. Terrestrial Manual: Avian influenza (infection with avian influenza viruses). 2019 26/11/2019].

3. Alexander, D.J., An overview of the epidemiology of avian influenza. Vaccine, 2007. 25(30): p. 5637–44.

4. European Commission, Council directive 2005/94/EC of 20 December 2005 on community measures for the control of avian influenza and repealing directive 92/40/EEC. Official Journal of the European Union, 2006. L10: 20/16.

5. OIE. Avian influenza. In: World Health Organization for Animal Health, Terrestrial Animal Health Code, 2017. Paris: OIE, chapter 10.4.. 2017 8 March 2022]; Available from: http://www.oie.int/fileadmin/Home/eng/Health_standards/tahc/current/chapitre_avian_influenza_viruses.pdf.

6. Vapnek, J., Regulatory measures against outbreaks of highly pathogenic avian influenza.. FAO Legal Papers Online #82, 2010.

7. Boni, M.F., et al., Economic epidemiology of avian influenza on smallholder poultry farms. Theor Popul Biol, 2013. 90: p. 135–44.

8. Sharp, G.B., et al., Coinfection of wild ducks by influenza A viruses: distribution patterns and biological significance. J Virol, 1997. 71(8): p. 6128–35.

9. Floyd, T., et al., Encephalitis and Death in Wild Mammals at a Rehabilitation Center after Infection with Highly Pathogenic Avian Influenza A(H5N8) Virus, United Kingdom. Emerg Infect Dis, 2021. 27(11): p. 2856–2863.

10. Adlhoch, C., et al., Avian influenza overview March - June 2022. Efsa j, 2022. 20(64): p. e07415.

11. Postel, A., et al., Infections with highly pathogenic avian influenza A virus (HPAIV) H5N8 in harbor seals at the German North Sea coast, 2021. Emerging Microbes & Infections, 2022. 11(1): p. 725–729.

12. Puryear, W., et al., Outbreak of Highly Pathogenic Avian Influenza H5N1 in New England Seals. bioRxiv, 2022:p. 2022.07.29.501155.

13. Rijks, J.M., et al., Highly Pathogenic Avian Influenza A(H5N1) Virus in Wild Red Foxes, the Netherlands, 2021. Emerg Infect Dis, 2021. 27(11): p. 2960–2962.

14. Eisfeld, A.J., G. Neumann, and Y. Kawaoka, At the centre: influenza A virus ribonucleoproteins. Nat Rev Microbiol, 2015. 13(1): p. 28–41.

15. Pauly, M.D., M.C. Procario, and A.S. Lauring, A novel twelve class fluctuation test reveals higher than expected mutation rates for influenza A viruses. eLife, 2017. 6: p. e26437.

16. Suárez, P., J. Valcárcel, and J. Ortín, Heterogeneity of the mutation rates of influenza A viruses: isolation of mutator mutants. J Virol, 1992. 66(4): p. 2491–4.

17. Suárez-López, P. and J. Ortín, An estimation of the nucleotide substitution rate at defined positions in the influenza virus haemagglutinin gene. J Gen Virol, 1994. 75 (Pt 2): p. 389–93.

18. Adlhoch, C., et al., Avian influenza overview February - May 2021. Efsa j, 2021. 19(12): p. e06951.

19. Adlhoch, C., et al., Avian influenza overview May - September 2021. Efsa j, 2022. 20(1): p. e07122.

20. Adlhoch, C., et al., Avian influenza overview December 2021 - March 2022. Efsa j, 2022. 20(4): p. e07289.

21. Pohlmann, A., et al., Has Epizootic Become Enzootic? Evidence for a Fundamental Change in the Infection Dynamics of Highly Pathogenic Avian Influenza in Europe, 2021. mBio, 2022.

22. Banyard, A.C., et al., Detection of Highly Pathogenic Avian Influenza Virus H5N1 Clade 2.3.4.4b in Great Skuas: A Species of Conservation Concern in Great Britain. Viruses, 2022. 14(2).

23. Caliendo, V., et al., Transatlantic spread of highly pathogenic avian influenza H5N1 by wild birds from Europe to North America in 2021. Scientific Reports, 2022. 12(1): p. 11729.

24. Bevins, S., et al., Intercontinental Movement of Highly Pathogenic Avian Influenza A(H5N1) Clade 2.3.4.4 Virus to the United States, 2021. Emerging Infectious Disease journal, 2022. 28(5): p. 1006.

25. Slomka, M.J., et al., Validated RealTime reverse transcriptase PCR methods for the diagnosis and pathotyping of Eurasian H7 avian influenza viruses. Influenza Other Respir Viruses, 2009. 3(4): p. 151–64.

26. Oliver, I., et al., A case of avian influenza A(H5N1) in England, January 2022. Euro Surveill, 2022. 27(5).

27. Hadfield, J., et al., Nextstrain: real-time tracking of pathogen evolution. Bioinformatics, 2018. 34(23): p. 4121–4123.

28. Katoh, K. and D.M. Standley, MAFFT multiple sequence alignment software version 7: improvements in performance and usability. Mol Biol Evol, 2013. 30(4): p. 772–80.

29. Larsson, A., AliView: a fast and lightweight alignment viewer and editor for large datasets. Bioinformatics, 2014. 30(22): p. 3276–3278.

30. Minh, B.Q., et al., IQ-TREE 2: New Models and Efficient Methods for Phylogenetic Inference in the Genomic Era. Molecular Biology and Evolution, 2020. 37(5): p. 1530–1534.

31. Kalyaanamoorthy, S., et al., ModelFinder: fast model selection for accurate phylogenetic estimates. Nature Methods, 2017. 14(6): p. 587–589.

32. Hoang, D.T., et al., UFBoot2: Improving the Ultrafast Bootstrap Approximation. Molecular Biology and Evolution, 2017. 35(2): p. 518–522.

33. Sagulenko, P., V. Puller, and R.A. Neher, TreeTime: Maximum-likelihood phylodynamic analysis. Virus Evolution, 2018. 4(1).

34. Yu, G., et al., ggtree: an r package for visualization and annotation of phylogenetic trees with their covariates and other associated data. Methods in Ecology and Evolution, 2017. 8(1): p. 28–36.

35. Wang, L.-G., et al., Treeio: An R Package for Phylogenetic Tree Input and Output with Richly Annotated and Associated Data. Molecular Biology and Evolution, 2019. 37(2): p. 599–603.

36. Poen, M.J., et al., Co-circulation of genetically distinct highly pathogenic avian influenza A clade 2.3.4.4 (H5N6) viruses in wild waterfowl and poultry in Europe and East Asia, 2017–18. Virus Evolution, 2019. 5(1).

37. CDC. H5N1 Genetic Changes Inventory: A Tool for Influenza Surveillance and Prepredness. 2012 25/07/2022]; Available from: https://www.cdc.gov/flu/pdf/avianflu/h5n1-inventory.pdf.

38. Suttie, A., et al., Inventory of molecular markers affecting biological characteristics of avian influenza A viruses. Virus Genes, 2019. 55(6): p. 739–768.

39. Suchard, M.A., et al., Bayesian phylogenetic and phylodynamic data integration using BEAST 1.10. Virus Evol, 2018. 4(1): p. vey016.

40. Ayres, D.L., et al., BEAGLE: An Application Programming Interface and High-Performance Computing Library for Statistical Phylogenetics. Systematic Biology, 2011. 61(1): p. 170–173.

41. Lemey, P., et al., Bayesian phylogeography finds its roots. PLoS Comput Biol, 2009. 5(9): p. e1000520.

42. Bielejec, F., et al., SpreaD3: Interactive Visualization of Spatiotemporal History and Trait Evolutionary Processes. Mol Biol Evol, 2016. 33(8): p. 2167–9.

43. Morey, R.D., J.-W. Romeijn, and J.N. Rouder, The philosophy of Bayes factors and the quantification of statistical evidence. Journal of Mathematical Psychology, 2016. 72: p. 6–18.

44. Bui, C.M., et al., Characterising routes of H5N1 and H7N9 spread in China using Bayesian phylogeographical analysis. Emerging Microbes & Infections, 2018. 7(1): p. 1–8.

45. Engelsma, M., et al., Multiple Introductions of Reassorted Highly Pathogenic Avian Influenza H5Nx Viruses Clade 2.3.4.4b Causing Outbreaks in Wild Birds and Poultry in The Netherlands, 2020-2021. Microbiology Spectrum, 2022. 10(2): p. e02499–21.

46. Duff, P., et al., Investigations associated with the 2020/21 highly pathogenic avian influenza epizootic in wild birds in Great Britain. Veterinary Record, 2021. 189(9): p. 356–358.

47. OFFLU. Influenza A Cleavage Sites. 2020 04/11/2022]; Available from: https://www.offlu.org/wp-content/uploads/2021/01/Influenza_A_Cleavage_Sites.pdf.

48. Wang, W., et al., Glycosylation at 158N of the hemagglutinin protein and receptor binding specificity synergistically affect the antigenicity and immunogenicity of a live attenuated H5N1 A/Vietnam/1203/2004 vaccine virus in ferrets. J Virol, 2010. 84(13): p. 6570–7.

49. Gao, Y., et al., Identification of amino acids in HA and PB2 critical for the transmission of H5N1 avian influenza viruses in a mammalian host. PLoS Pathog, 2009. 5(12): p. e1000709.

50. Abed, Y., N. Goyette, and G. Boivin, Generation and characterization of recombinant influenza A (H1N1) viruses harboring amantadine resistance mutations. Antimicrob Agents Chemother, 2005. 49(2): p. 556–9.

51. Bean, W.J., S.C. Threlkeld, and R.G. Webster, Biologic potential of amantadine-resistant influenza A virus in an avian model. J Infect Dis, 1989. 159(6): p. 1050–6.

52. Cheung, C.L., et al., Distribution of amantadine-resistant H5N1 avian influenza variants in Asia. J Infect Dis, 2006. 193(12): p. 1626–9.

53. Ilyushina, N.A., E.A. Govorkova, and R.G. Webster, Detection of amantadine-resistant variants among avian influenza viruses isolated in North America and Asia. Virology, 2005. 341(1): p. 102–6.

54. Lan, Y., et al., A comprehensive surveillance of adamantane resistance among human influenza A virus isolated from mainland China between 1956 and 2009. Antivir Ther, 2010. 15(6): p. 853–9.

55. Kode, S.S., et al., A novel I117T substitution in neuraminidase of highly pathogenic avian influenza H5N1 virus conferring reduced susceptibility to oseltamivir and zanamivir. Vet Microbiol, 2019. 235: p. 21–24.

56. Omoto, S., et al., Characterization of influenza virus variants induced by treatment with the endonuclease inhibitor baloxavir marboxil. Sci Rep, 2018. 8(1): p. 9633.

57. APHA. Livestock Demographic Data Group: Poultry Population Report. 2019 16/09/2022]; Available from: http://apha.defra.gov.uk/documents/surveillance/diseases/lddg-pop-report-avian2019.pdf.

58. Lewis, N.S., et al., Emergence and spread of novel H5N8, H5N5 and H5N1 clade 2.3.4.4 highly pathogenic avian influenza in 2020. Emerging Microbes & Infections, 2021. 10(1): p. 148–151.

59. King, J., et al., Highly pathogenic avian influenza virus incursions of subtype H5N8, H5N5, H5N1, H5N4, and H5N3 in Germany during 2020-21. Virus Evolution, 2022. 8(1).

60. Cheng, K., et al., PB2-E627K and PA-T97I substitutions enhance polymerase activity and confer a virulent phenotype to an H6N1 avian influenza virus in mice. Virology, 2014. 468-470: p. 207–213.

61. de Jong, R.M., et al., Rapid emergence of a virulent PB2 E627K variant during adaptation of highly pathogenic avian influenza H7N7 virus to mice. Virol J, 2013. 10: p. 276.

62. Hatta, M., et al., Growth of H5N1 influenza A viruses in the upper respiratory tracts of mice. PLoS Pathog, 2007. 3(10): p. 1374–9.

63. Herfst, S., et al., Airborne transmission of influenza A/H5N1 virus between ferrets. Science, 2012. 336(6088): p. 1534–41.

64. Sediri, H., et al., PB2 subunit of avian influenza virus subtype H9N2: a pandemic risk factor. J Gen Virol, 2016. 97(1): p. 39–48.

65. Liu, H., et al., Phylogenetic and Phylogeographic Analysis of the Highly Pathogenic H5N6 Avian Influenza Virus in China. Viruses, 2022. 14(8).

66. Htwe, K.T.Z., et al., Phylogeographic analysis of human influenza A and B viruses in Myanmar, 2010–2015. PLOS ONE, 2019. 14(1): p. e0210550.

67. Duchatel, F., B.M.d.C. Bronsvoort, and S. Lycett, Phylogeographic Analysis and Identification of Factors Impacting the Diffusion of Foot-and-Mouth Disease Virus in Africa. Frontiers in Ecology and Evolution, 2019. 7.

68. Moreira Salles, A.P., et al., Updating the Phylodynamics of Yellow Fever Virus 2016–2019 Brazilian Outbreak With New 2018 and 2019 São Paulo Genomes. Frontiers in Microbiology, 2022. 13.

