## Supplemental Figures for "Investigating the genetic diversity of H5 avian influenza in the UK 2020-2022"

**Figure S1.** Time-resolved maximum-likelihood phylogenetic trees containing the H5Nx sequences obtained from the UK, with relevant global reference sequences. (A) H5, (B) NA, (C) PB2, (D) PB1, (E) PA, (F) NP, (G) MP and (H) NS. Sequences are coloured according the H5 subtype, and UK H5Nx genotypes are illustrated. The sequences obtained from the UK are indicated with circular tip shapes.

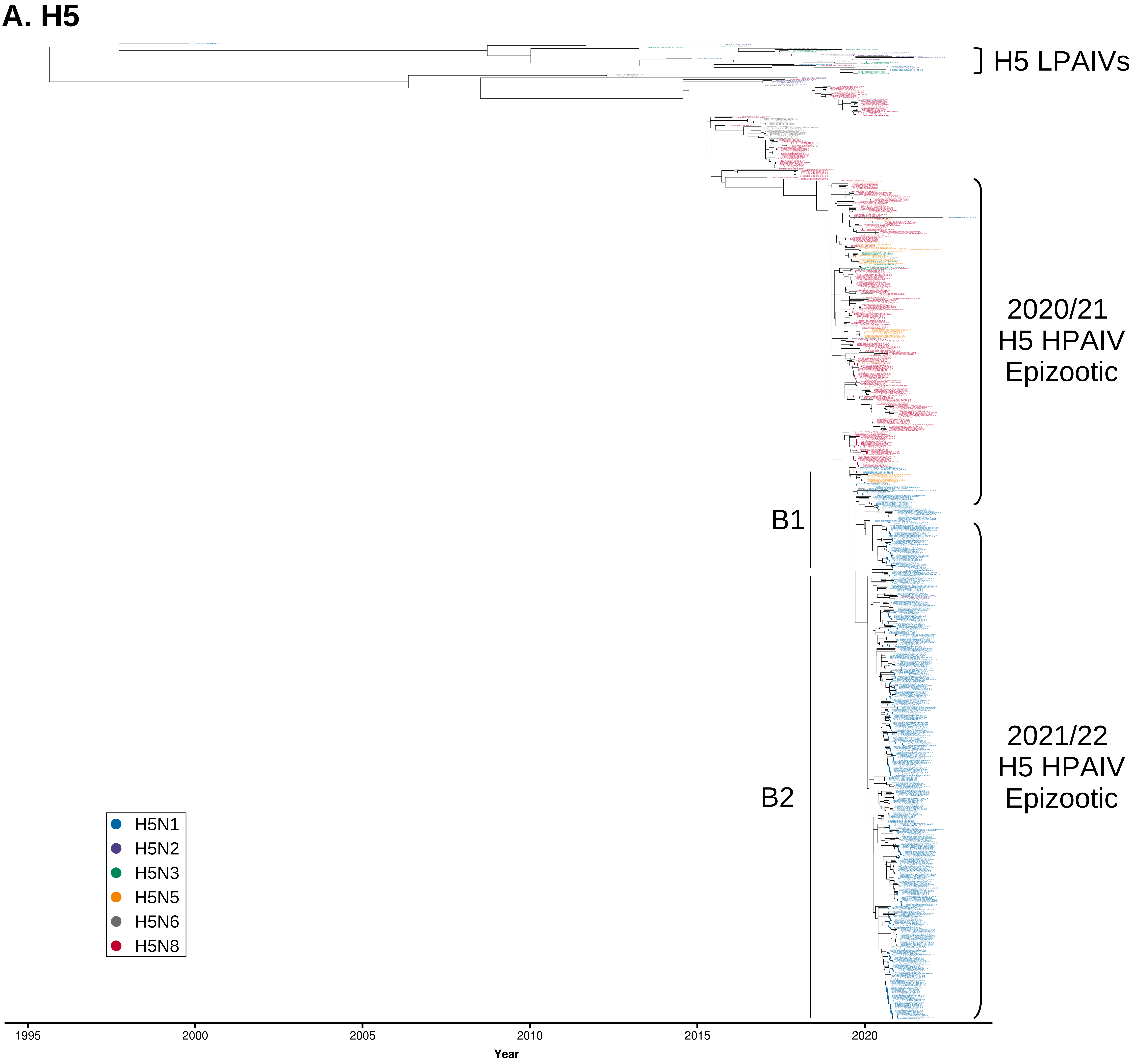
A. H5
H5 LPAIVs
2020/21
H5 HPAIV
Epizootic
2021/22
H5 HPAIV
Epizootic
B1
B2
H5N1
H5N2
H5N3
H5N5
H5N6
H5N8
1995
2000
2005
2010
2015
2020
Year

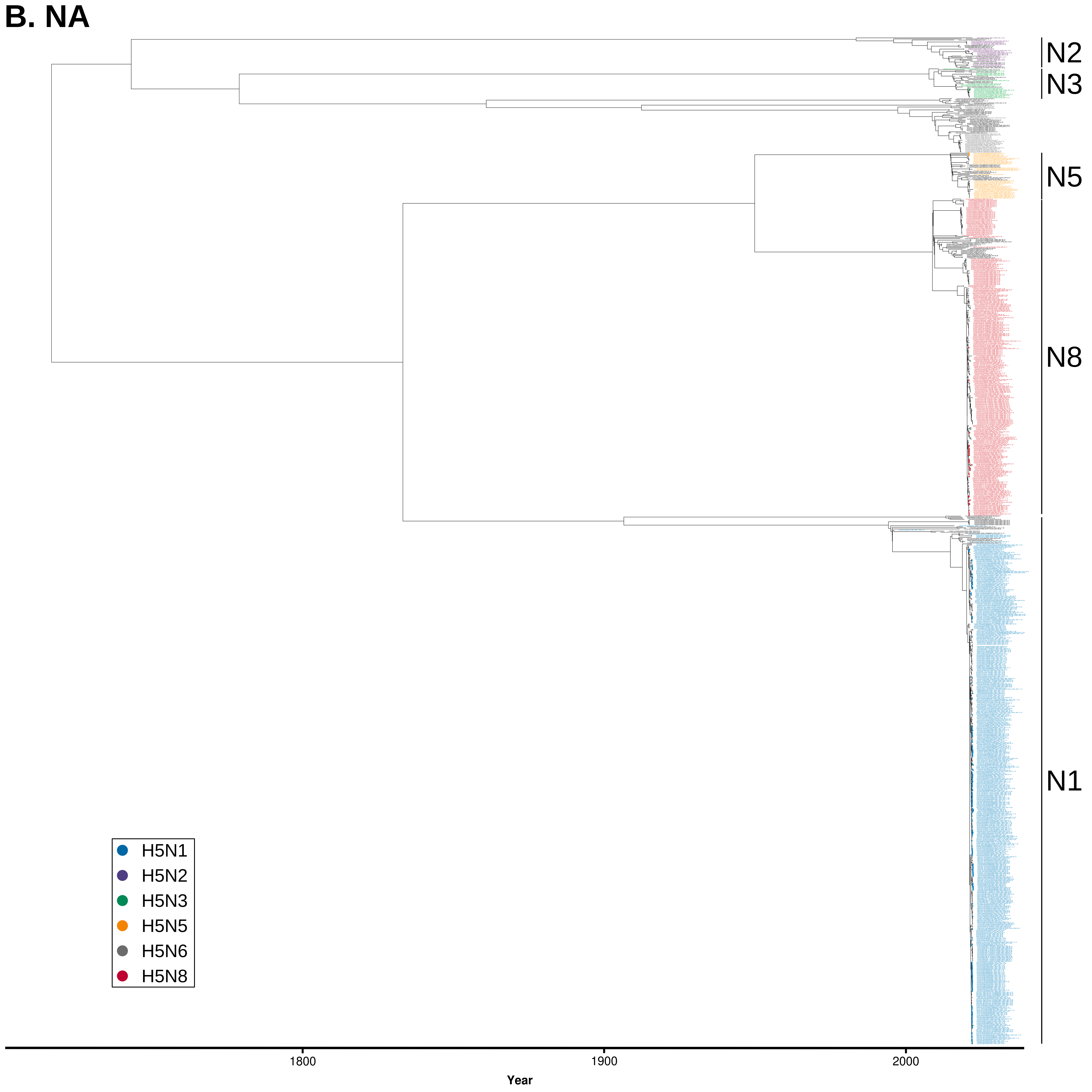
B. NA
N2
N3
N5
N8
N1
H5N1
H5N2
H5N3
H5N5
H5N6
H5N8
1800
1900
2000
Year

# C. PB2

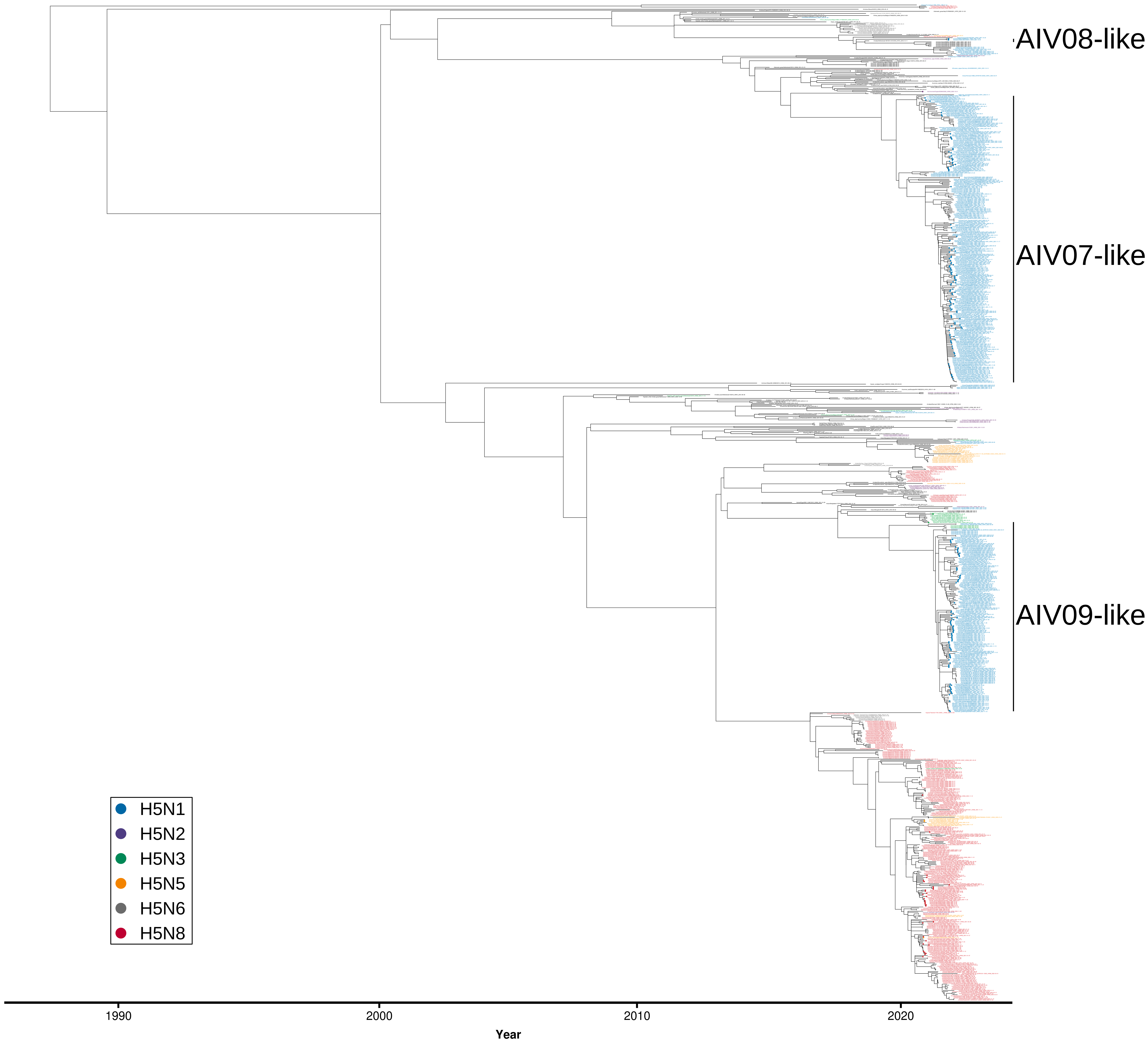
AIV08-like
AIV07-like
AIV09-like
H5N1
H5N2
H5N3
H5N5
H5N6
H5N8
1990
2000
2010
2020
Year

# D. PB1

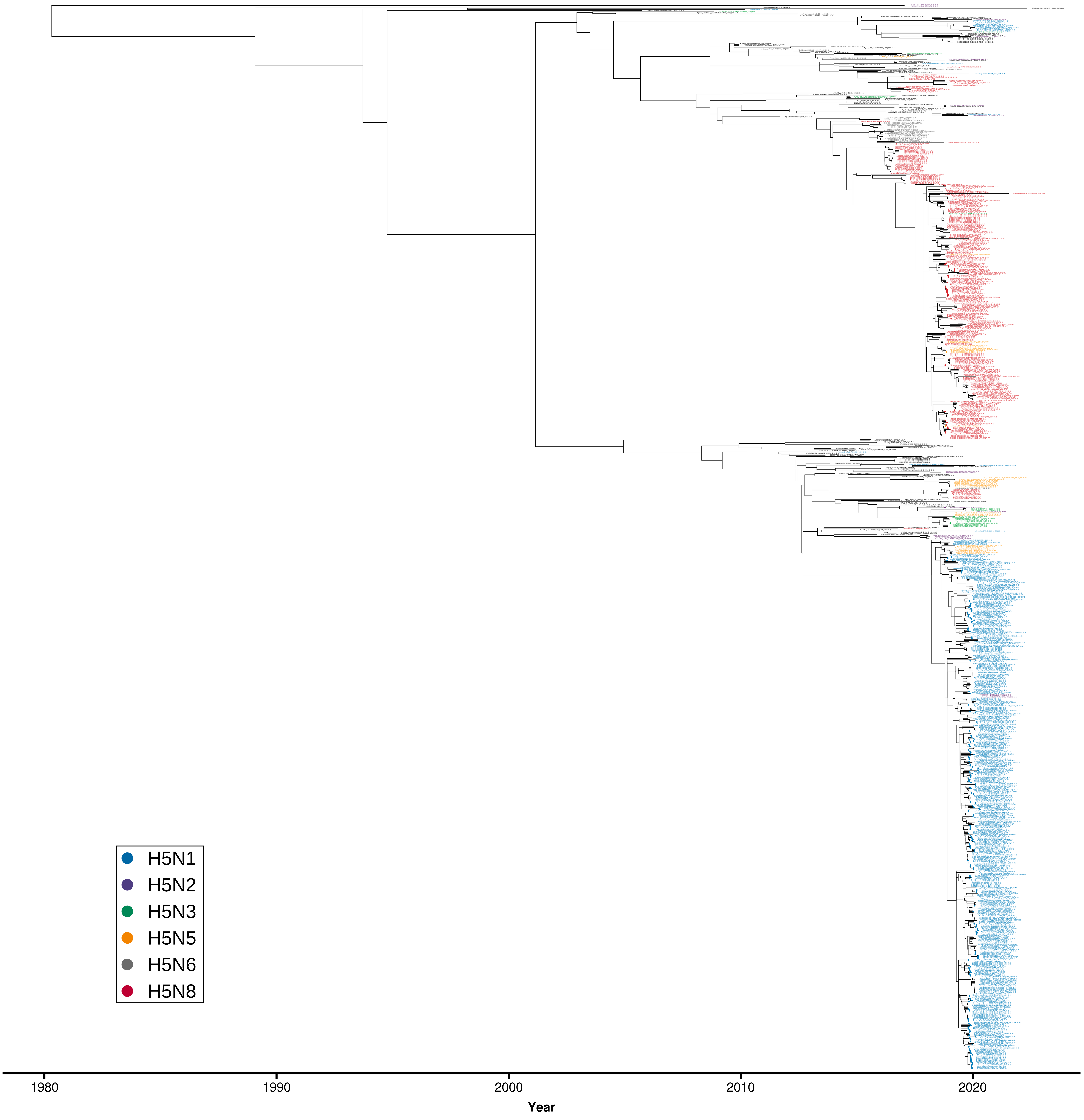

E. PA

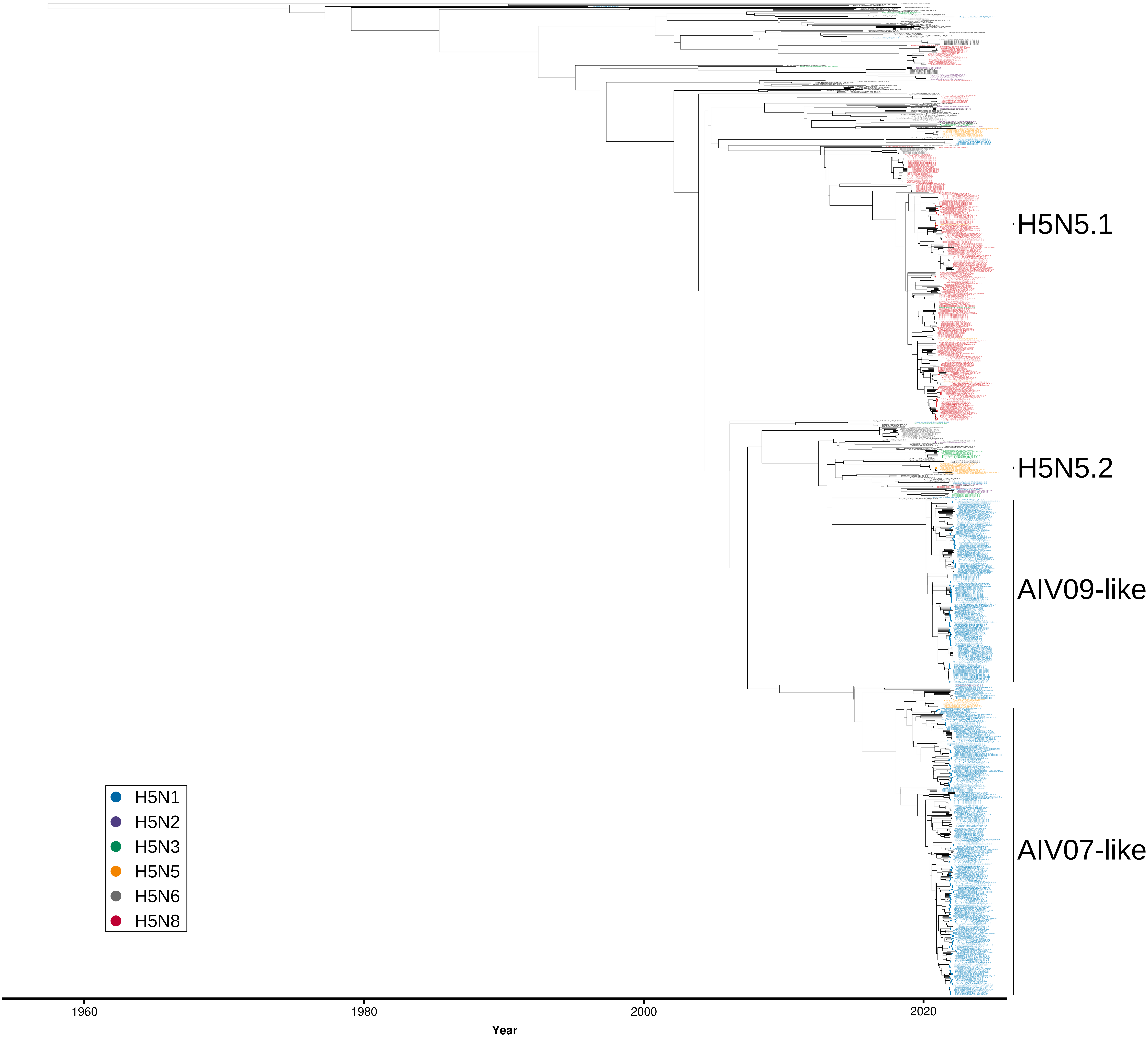
H5N5.1
H5N5.2
AIV09-like
AIV07-like
H5N1
H5N2
H5N3
H5N5
H5N6
H5N8
1960
1980
2000
2020
Year

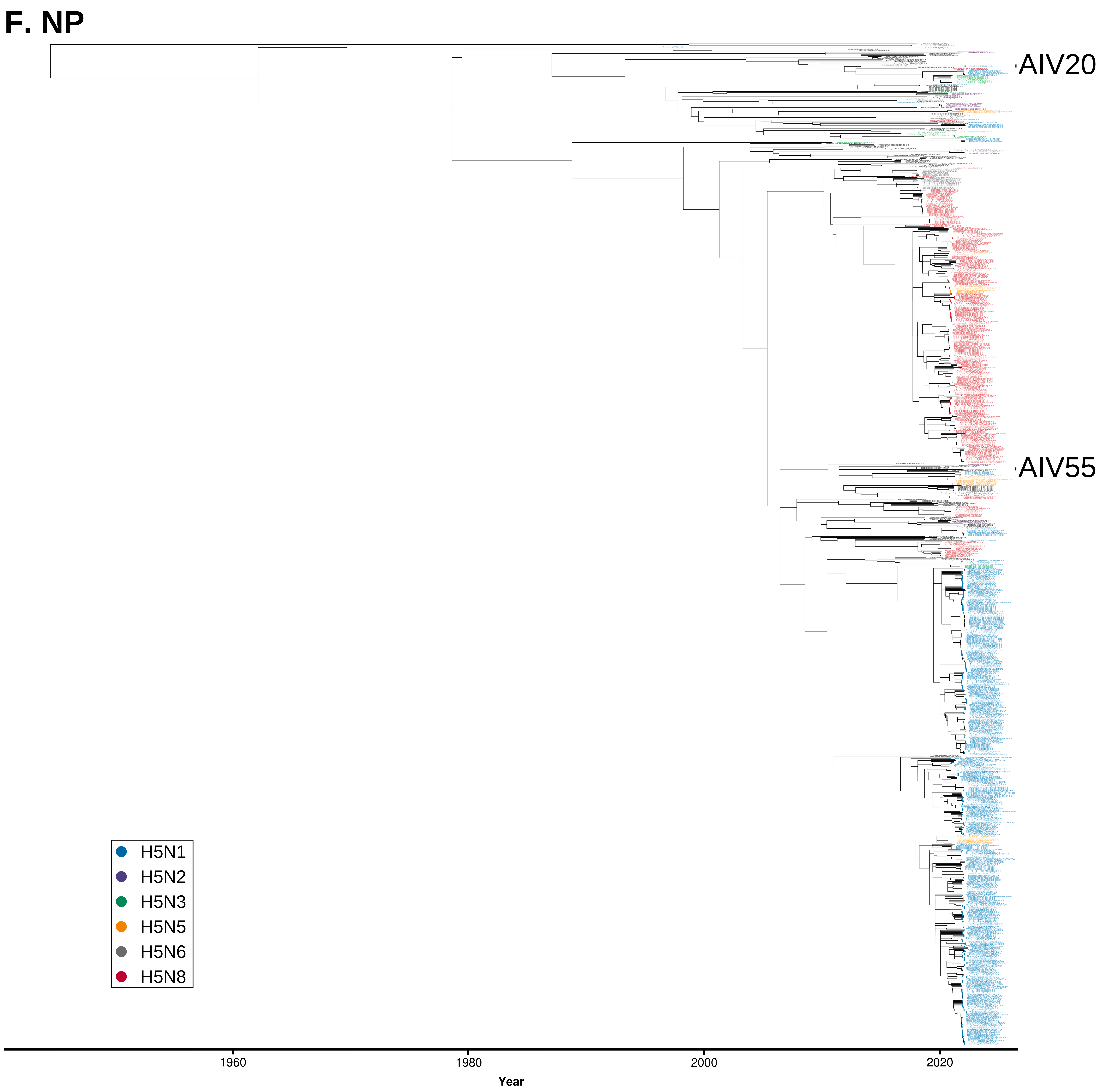
F. NP
AIV20
AIV55
H5N1
H5N2
H5N3
H5N5
H5N6
H5N8
1960
1980
2000
2020
Year

# G. MP

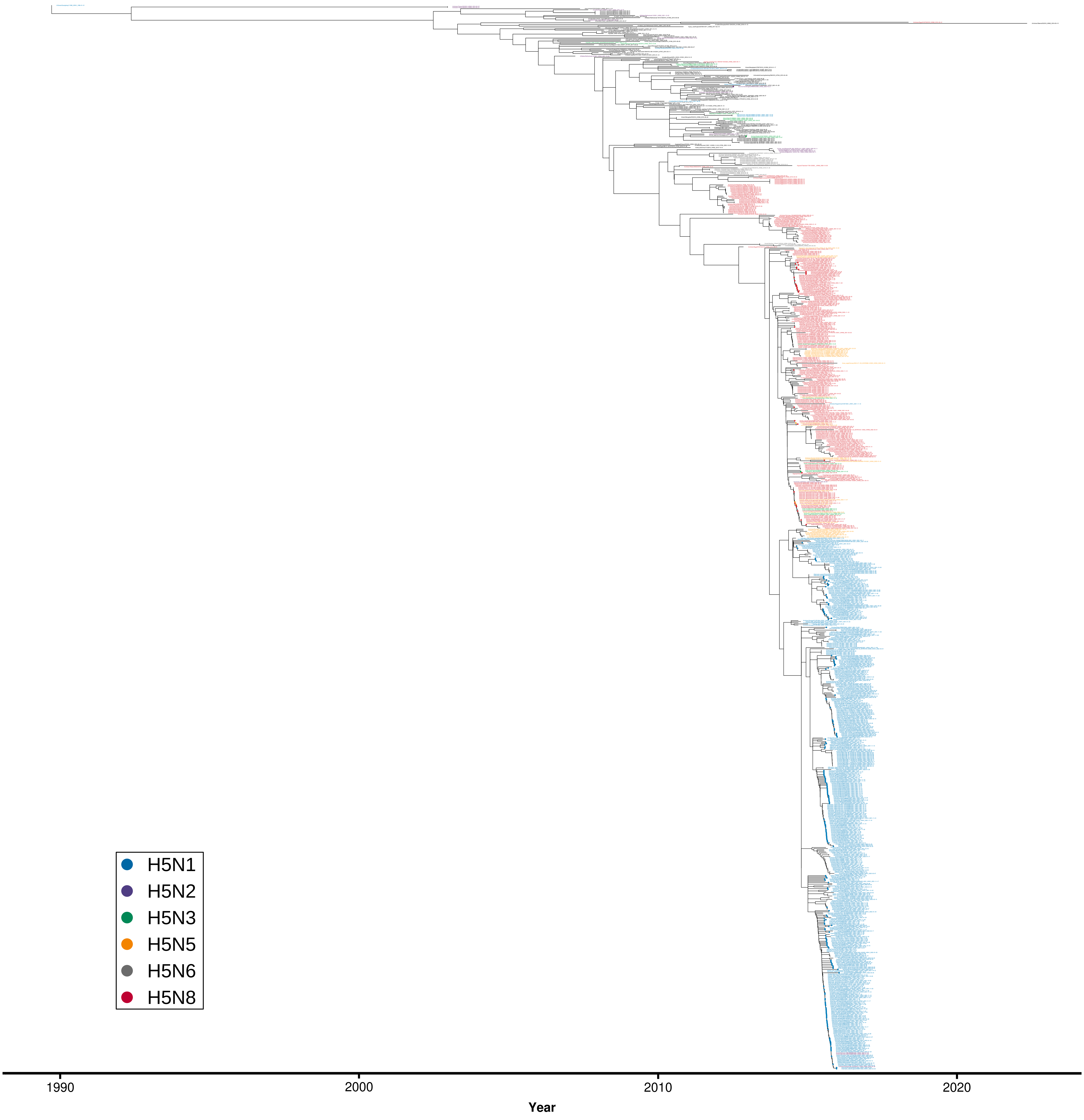

# H. NS

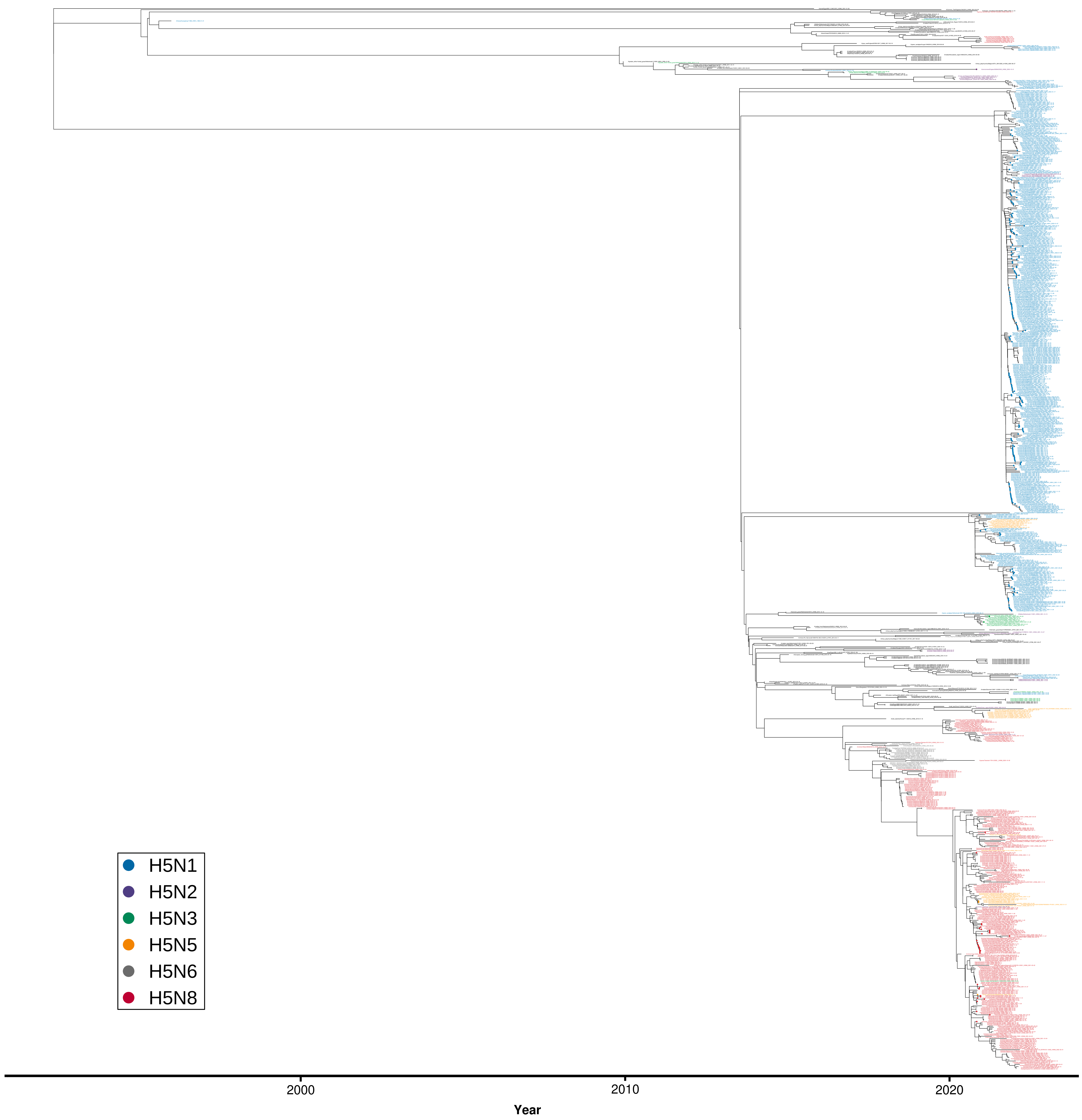

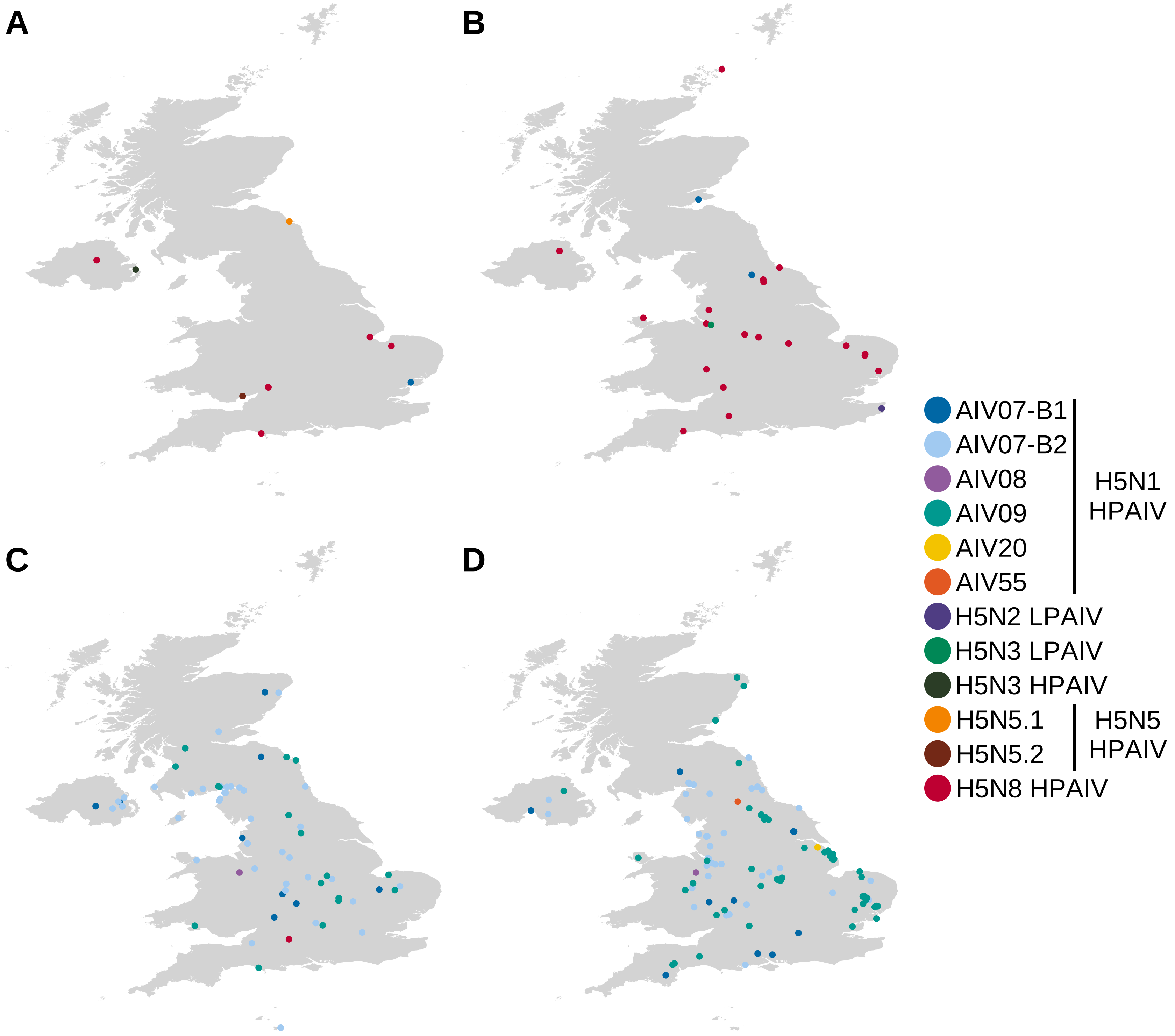


**Figure S2.** Geographic distribution of H5Nx AIVs that were sequenced from wild birds (A and C) and poultry (B and D) during the 2020/21 (A and B) and 2021/22 (C and D) epizootics in the UK. Locations are coloured according to AIV subtype, pathotype and genotype.
