## Supplemental Tables for "Investigating the genetic diversity of H5 avian influenza in the UK 2020-2022"

**Table S1.** GISAID EpiFlu accession numbers for all sequences generated in this study.

| Accession Number | Isolate Name | Subtype | Collection Date | Genotype |
|---|---|---|---|---|
| EPI_ISL_710508 | A/Greylag goose/England/033100/2020 | H5N8 | 2020-10-30 | H5N8 |
| EPI_ISL_13243697 | A/environment/England/030642/2020 | H5N2 | 2020-10-31 | H5N2 LPAIV |
| EPI_ISL_626652 | A/chicken/England/030720/2020 | H5N8 | 2020-11-02 | H5N8 |
| EPI_ISL_710506 | A/Canada goose/England/032697/2020 | H5N8 | 2020-11-03 | H5N8 |
| EPI_ISL_710507 | A/Greylag goose/England/032698/2020 | H5N8 | 2020-11-03 | H5N8 |
| EPI_ISL_1123357 | A/Brent goose/England/233339/2020 | H5N8 | 2020-11-08 | H5N8 |
| EPI_ISL_710509 | A/chicken/England/033708/2020 | H5N8 | 2020-11-10 | H5N8 |
| EPI_ISL_1123359 | A/mute swan/England/263814/2020 | H5N8 | 2020-11-10 | H5N8 |
| EPI_ISL_1123358 | A/Brent goose/England/095684/2020 | H5N5 | 2020-11-12 | H5N5.1 |
| EPI_ISL_11402745 | A/mute swan/Northern Ireland/VMDL20-301/2020 | H5N8 | 2020-11-12 | H5N8 |
| EPI_ISL_710512 | A/whistling duck/England/035643/2020 | H5N8 | 2020-11-19 | H5N8 |
| EPI_ISL_710511 | A/chicken/England/037052/2020 | H5N8 | 2020-11-21 | H5N8 |
| EPI_ISL_683997 | A/mute swan/Wales/048069/2020 | H5N5 | 2020-11-24 | H5N5.2 |
| EPI_ISL_683999 | A/mute swan/Wales/048068/2020 | H5N5 | 2020-11-24 | H5N5.2 |
| EPI_ISL_710504 | A/turkey/England/037784/2020 | H5N8 | 2020-11-28 | H5N8 |
| EPI_ISL_1123360 | A/mute swan/England/234135/2020 | H5N8 | 2020-12-01 | H5N8 |
| EPI_ISL_710505 | A/turkey/England/038115/2020 | H5N8 | 2020-12-02 | H5N8 |
| EPI_ISL_766876 | A/mute swan/England/234255/2020 | H5N1 | 2020-12-03 | AIV07-B1 |
| EPI_ISL_766052 | A/turkey/England/038730/2020 | H5N8 | 2020-12-03 | H5N8 |
| EPI_ISL_766053 | A/turkey/England/039352/2020 | H5N8 | 2020-12-04 | H5N8 |
| EPI_ISL_766054 | A/turkey/England/039472/2020 | H5N8 | 2020-12-04 | H5N8 |
| EPI_ISL_2081527 | A/red fox/England/AVP-M1-21-01/2020 | H5N8 | 2020-12-08 | H5N8 |
| EPI_ISL_2081528 | A/seal/England/AVP-031141/2020 | H5N8 | 2020-12-08 | H5N8 |
| EPI_ISL_766056 | A/falcon/England/041976/2020 | H5N8 | 2020-12-14 | H5N8 |
| EPI_ISL_1122425 | A/chicken/England/043315/2020 | H5N1 | 2020-12-15 | AIV07-B1 |
| EPI_ISL_1123263 | A/chicken/Scotland/043405/2020 | H5N8 | 2020-12-16 | H5N8 |
| EPI_ISL_1123350 | A/chicken/England/043683/2020 | H5N8 | 2020-12-17 | H5N8 |
| EPI_ISL_1123351 | A/duck/England/043628/2020 | H5N8 | 2020-12-18 | H5N8 |
| EPI_ISL_1123352 | A/chicken/England/045984/2020 | H5N8 | 2020-12-25 | H5N8 |
| EPI_ISL_1123353 | A/duck/England/046311/2020 | H5N8 | 2020-12-27 | H5N8 |
| EPI_ISL_1123354 | A/chicken/England/046491/2020 | H5N8 | 2020-12-28 | H5N8 |
| EPI_ISL_11402752 | A/chicken/Northern Ireland/029599/2020 | H5N8 | 2020-12-31 | H5N8 |
| EPI_ISL_11392591 | A/peregrine falcon/Northern Ireland/AI102021-2/2021 | H5N3 | 2021-01-22 | H5N3 HPAIV |
| EPI_ISL_1123355 | A/pheasant/Wales/000252/2021 | H5N8 | 2021-01-27 | H5N8 |
| EPI_ISL_11402307 | A/chicken/England/010821/2021 | H5N8 | 2021-02-06 | H5N8 |
| EPI_ISL_1123361 | A/pheasant/Scotland/000348/2021 | H5N1 | 2021-02-10 | AIV07-B1 |
| EPI_ISL_11402318 | A/turkey/England/018179/2021 | H5N3 | 2021-03-25 | H5N3 LPAIV |
| EPI_ISL_11406397 | A/chicken/England/019018/2021 | H5N8 | 2021-03-28 | H5N8 |
| EPI_ISL_11406400 | A/raven/England/019024/2021 | H5N8 | 2021-03-28 | H5N8 |
| EPI_ISL_11406449 | A/pheasant/England/019028/2021 | H5N8 | 2021-03-28 | H5N8 |
| EPI_ISL_11388317 | A/falcon/England/AVP-21-019051A/2021 | H5N8 | 2021-03-30 | H5N8 |
| EPI_ISL_5804708 | A/mute swan/England/053054/2021 | H5N1 | 2021-10-24 | AIV07-B1 |
| EPI_ISL_9012457 | A/chicken/England/053052/2021 | H5N1 | 2021-10-24 | AIV07-B1 |
| EPI_ISL_9012572 | A/pheasant/Wales/385129/2021 | H5N1 | 2021-10-27 | AIV08 |
| EPI_ISL_9012618 | A/chicken/Wales/053969/2021 | H5N1 | 2021-10-30 | AIV08 |
| EPI_ISL_9029965 | A/Greylag goose/England/054503/2021 | H5N1 | 2021-10-30 | AIV07-B2 |
| EPI_ISL_9012696 | A/chicken/Scotland/054477/2021 | H5N1 | 2021-11-01 | AIV09 |
| EPI_ISL_9029961 | A/Canada goose/England/385250/2021 | H5N1 | 2021-11-01 | AIV07-B1 |
| EPI_ISL_90122694 | A/guineafowl/Scotland/054471/2021 | H5N1 | 2021-11-01 | AIV09 |
| EPI_ISL_9012700 | A/domestic duck/Scotland/054469/2021 | H5N1 | 2021-11-01 | AIV09 |
| EPI_ISL_13369398 | A/buzzard/England/107870/2021 | H5N1 | 2021-11-02 | AIV09 |
| EPI_ISL_13369399 | A/pheasant/England/107851/2021 | H5N1 | 2021-11-02 | AIV07-B1 |
| EPI_ISL_13369400 | A/Herring gull/Wales/054692/2021 | H5N1 | 2021-11-02 | AIV09 |
| EPI_ISL_13369401 | A/mute swan/England/243602/2021 | H5N1 | 2021-11-03 | AIV09 |

### Table S1 continued.

| Accession Number | Isolate Name | Subtype | Collection Date | Genotype |
|---|---|---|---|---|
| EPI_ISL_13369741 | A/whooper swan/England/055405/2021 | H5N1 | 2021-11-05 | AIV07-B2 |
| EPI_ISL_8814146 | A/turkey/England/055251/2021 | H5N1 | 2021-11-06 | AIV07-B2 |
| EPI_ISL_13369742 | A/mute swan/England/298902/2021 | H5N8 | 2021-11-07 | H5N8 |
| EPI_ISL_13369743 | A/mallard/England/056080/2021 | H5N1 | 2021-11-08 | AIV07-B2 |
| EPI_ISL_13369744 | A/barnacle goose/England/292151/2021 | H5N1 | 2021-11-08 | AIV07-B2 |
| EPI_ISL_9029962 | A/Whooper swan/Scotland/056219/2021 | H5N1 | 2021-11-09 | AIV07-B2 |
| EPI_ISL_13369745 | A/chicken/England/056017/2021 | H5N1 | 2021-11-10 | AIV09 |
| EPI_ISL_8814195 | A/turkey/England/056764/2021 | H5N1 | 2021-11-10 | AIV07-B2 |
| EPI_ISL_9029960 | A/mute swan/England/385466/2021 | H5N1 | 2021-11-11 | AIV07-B1 |
| EPI_ISL_13369746 | A/chicken/England/057016/2021 | H5N1 | 2021-11-12 | AIV09 |
| EPI_ISL_13369747 | A/Barnacle goose/England/108089/2021 | H5N1 | 2021-11-12 | AIV09 |
| EPI_ISL_13369748 | A/mute swan/England/108094/2021 | H5N1 | 2021-11-12 | AIV09 |
| EPI_ISL_13369753 | A/barnacle goose/Scotland/292583/2021 | H5N1 | 2021-11-12 | AIV09 |
| EPI_ISL_13369754 | A/mute swan/England/243983/2021 | H5N1 | 2021-11-13 | AIV09 |
| EPI_ISL_13369755 | A/Pink-footed goose/England/243989/2021 | H5N1 | 2021-11-13 | AIV09 |
| EPI_ISL_13369756 | A/mute swan/England/244029/2021 | H5N1 | 2021-11-14 | AIV07-B2 |
| EPI_ISL_13369757 | A/mallard/Scotland/057554/2021 | H5N1 | 2021-11-14 | AIV07-B2 |
| EPI_ISL_9029956 | A/chicken/England/057314/2021 | H5N1 | 2021-11-14 | AIV07-B2 |
| EPI_ISL_13369758 | A/mute swan/England/292632/2021 | H5N1 | 2021-11-15 | AIV07-B1 |
| EPI_ISL_13369759 | A/chicken/England/058797/2021 | H5N1 | 2021-11-16 | AIV07-B2 |
| EPI_ISL_13369760 | A/pink-footed goose/Scotland/058488/2021 | H5N1 | 2021-11-16 | AIV07-B2 |
| EPI_ISL_13369761 | A/Kestrel/Scotland/058512/2021 | H5N1 | 2021-11-16 | AIV07-B2 |
| EPI_ISL_9029957 | A/turkey/England/057679/2021 | H5N1 | 2021-11-16 | AIV07-B2 |
| EPI_ISL_13369762 | A/barnacle goose/England/061307/2021 | H5N1 | 2021-11-17 | AIV09 |
| EPI_ISL_9029959 | A/domestic duck/England/058612/2021 | H5N1 | 2021-11-18 | AIV07-B2 |
| EPI_ISL_13369763 | A/turkey/England/059231/2021 | H5N1 | 2021-11-19 | AIV09 |
| EPI_ISL_13369764 | A/domestic goose/England/059141/2021 | H5N1 | 2021-11-19 | AIV09 |
| EPI_ISL_13369765 | A/barnacle goose/Scotland/062003/2021 | H5N1 | 2021-11-19 | AIV07-B2 |
| EPI_ISL_13369766 | A/barnacle goose/Scotland/062007/2021 | H5N1 | 2021-11-19 | AIV09 |
| EPI_ISL_13369767 | A/turkey/England/059407/2021 | H5N1 | 2021-11-20 | AIV09 |
| EPI_ISL_13369768 | A/turkey/England/059498/2021 | H5N1 | 2021-11-20 | AIV09 |
| EPI_ISL_13369769 | A/chicken/England/59973/2021 | H5N1 | 2021-11-20 | AIV09 |
| EPI_ISL_13369770 | A/common buzzard/England/300284/2021 | H5N1 | 2021-11-20 | AIV07-B2 |
| EPI_ISL_13369771 | A/turkey/England/059870/2021 | H5N1 | 2021-11-21 | AIV09 |
| EPI_ISL_13369772 | A/chicken/England/061210/2021 | H5N1 | 2021-11-22 | AIV09 |
| EPI_ISL_13369773 | A/chicken/England/061529/2021 | H5N1 | 2021-11-22 | AIV07-B2 |
| EPI_ISL_13369774 | A/black-headed gull/England/386075/2021 | H5N1 | 2021-11-22 | AIV07-B2 |
| EPI_ISL_13369775 | A/peafowl/England/061723/2021 | H5N1 | 2021-11-23 | AIV07-B2 |
| EPI_ISL_13370248 | A/domestic duck/Wales/061914/2021 | H5N1 | 2021-11-23 | AIV09 |
| EPI_ISL_13369776 | A/greylag goose/Northern Ireland/17077/2021 | H5N1 | 2021-11-24 | AIV07-B1 |
| EPI_ISL_13369777 | A/chicken/England/062137/2021 | H5N1 | 2021-11-25 | AIV09 |
| EPI_ISL_13369778 | A/mute swan/Northern Ireland/17176/2021 | H5N1 | 2021-11-25 | AIV07-B2 |
| EPI_ISL_13369779 | A/barnacle goose/England/293824/2021 | H5N1 | 2021-11-26 | AIV07-B2 |
| EPI_ISL_13369780 | A/barnacle goose/England/293809/2021 | H5N1 | 2021-11-26 | AIV07-B2 |
| EPI_ISL_13369781 | A/chicken/Northern Ireland/17463/2021 | H5N1 | 2021-11-26 | AIV09 |
| EPI_ISL_13369782 | A/turkey/England/062369/2021 | H5N1 | 2021-11-27 | AIV09 |
| EPI_ISL_13369783 | A/Canada goose/England/300363/2021 | H5N1 | 2021-11-27 | AIV09 |
| EPI_ISL_13369784 | A/chicken/England/063664/2021 | H5N1 | 2021-11-28 | AIV09 |
| EPI_ISL_13370249 | A/domestic duck/Northern Ireland/17400/2021 | H5N1 | 2021-11-28 | AIV07-B1 |
| EPI_ISL_13369809 | A/greylag goose/England/108485/2021 | H5N1 | 2021-11-29 | AIV07-B2 |
| EPI_ISL_13370343 | A/buzzard/Wales/055388/2021 | H5N1 | 2021-11-29 | AIV07-B2 |
| EPI_ISL_13370415 | A/mute swan/Northern Ireland/17269/2021 | H5N1 | 2021-11-29 | AIV07-B2 |
| EPI_ISL_13370416 | A/Black-headed gull/England/064433/2021 | H5N1 | 2021-11-30 | AIV07-B2 |
| EPI_ISL_13370417 | A/barnacle goose/Scotland/064368/2021 | H5N1 | 2021-11-30 | AIV07-B2 |

**Table S1 continued.**

| Accession Number | Isolate Name | Subtype | Collection Date | Genotype |
|---|---|---|---|---|
| EPI_ISL_13370418 | A/common buzzard/Scotland/294149/2021 | H5N1 | 2021-11-30 | AIV07-B2 |
| EPI_ISL_13370419 | A/chicken/England/064357/2021 | H5N1 | 2021-12-01 | AIV09 |
| EPI_ISL_13370420 | A/chicken/England/064233/2021 | H5N1 | 2021-12-01 | AIV09 |
| EPI_ISL_13370421 | A/chicken/England/064131/2021 | H5N1 | 2021-12-01 | AIV07-B1 |
| EPI_ISL_13370505 | A/kestrel/England/244611/2021 | H5N1 | 2021-12-01 | AIV09 |
| EPI_ISL_13370506 | A/Mute Swan/England/294146/2021 | H5N1 | 2021-12-01 | AIV07-B2 |
| EPI_ISL_13370507 | A/turkey/England/064661/2021 | H5N1 | 2021-12-02 | AIV09 |
| EPI_ISL_13370508 | A/mute swan/England/244635/2021 | H5N1 | 2021-12-02 | AIV09 |
| EPI_ISL_13370509 | A/mute swan/Northern Ireland/17404/2021 | H5N1 | 2021-12-02 | AIV07-B2 |
| EPI_ISL_13370510 | A/chicken/Scotland/064936/2021 | H5N1 | 2021-12-02 | AIV07-B2 |
| EPI_ISL_13370511 | A/turkey/Wales/065047/2021 | H5N1 | 2021-12-02 | AIV07-B2 |
| EPI_ISL_13370512 | A/mute swan/Northern Ireland/17533/2021 | H5N1 | 2021-12-03 | AIV07-B1 |
| EPI_ISL_13370892 | A/Lapwing/England/067574/2021 | H5N1 | 2021-12-03 | AIV07-B2 |
| EPI_ISL_13370513 | A/domestic duck/England/065343/2021 | H5N1 | 2021-12-04 | AIV07-B2 |
| EPI_ISL_13370514 | A/chicken/England/065487/2021 | H5N1 | 2021-12-04 | AIV09 |
| EPI_ISL_13370515 | A/chicken/England/066019/2021 | H5N1 | 2021-12-04 | AIV09 |
| EPI_ISL_13370516 | A/Greylag goose/England/108622/2021 | H5N1 | 2021-12-04 | AIV07-B2 |
| EPI_ISL_13370517 | A/barnacle goose/England/294327/2021 | H5N1 | 2021-12-04 | AIV07-B2 |
| EPI_ISL_13370518 | A/chicken/England/066782/2021 | H5N1 | 2021-12-05 | AIV09 |
| EPI_ISL_13370519 | A/domestic duck/England/066456/2021 | H5N1 | 2021-12-06 | AIV07-B1 |
| EPI_ISL_13370520 | A/mute swan/England/244574/2021 | H5N1 | 2021-12-06 | AIV09 |
| EPI_ISL_13370521 | A/mute swan/Northern Ireland/17634/2021 | H5N1 | 2021-12-06 | AIV07-B2 |
| EPI_ISL_13370522 | A/barnacle goose/Scotland/072143/2021 | H5N1 | 2021-12-06 | AIV07-B2 |
| EPI_ISL_13370523 | A/chicken/England/067219/2021 | H5N1 | 2021-12-08 | AIV09 |
| EPI_ISL_13370524 | A/turkey/England/067429/2021 | H5N1 | 2021-12-08 | AIV09 |
| EPI_ISL_13370556 | A/greylag goose/Northern Ireland/17795/2021 | H5N1 | 2021-12-08 | AIV07-B2 |
| EPI_ISL_13370557 | A/chicken/Scotland/068163/2021 | H5N1 | 2021-12-08 | AIV07-B2 |
| EPI_ISL_13370558 | A/chicken/England/068375/2021 | H5N1 | 2021-12-09 | AIV09 |
| EPI_ISL_13370559 | A/turkey/England/068583/2021 | H5N1 | 2021-12-09 | AIV07-B2 |
| EPI_ISL_13370560 | A/pheasant/England/294628/2021 | H5N1 | 2021-12-09 | AIV07-B2 |
| EPI_ISL_13370561 | A/barnacle goose/England/294620/2021 | H5N1 | 2021-12-09 | AIV07-B2 |
| EPI_ISL_13370562 | A/chicken/Scotland/068270/2021 | H5N1 | 2021-12-09 | AIV07-B1 |
| EPI_ISL_13370563 | A/domestic duck/England/069098/2021 | H5N1 | 2021-12-10 | AIV07-B2 |
| EPI_ISL_13370564 | A/chicken/Northern Ireland/17903/2021 | H5N1 | 2021-12-10 | AIV07-B2 |
| EPI_ISL_13370568 | A/domestic duck/Northern Ireland/17961/2021 | H5N1 | 2021-12-10 | AIV07-B2 |
| EPI_ISL_13370569 | A/chicken/England/069549/2021 | H5N1 | 2021-12-11 | AIV09 |
| EPI_ISL_13370570 | A/chicken/England/070105/2021 | H5N1 | 2021-12-11 | AIV09 |
| EPI_ISL_13370571 | A/chicken/England/069816/2021 | H5N1 | 2021-12-12 | AIV55 |
| EPI_ISL_13370572 | A/domestic duck/England/070195/2021 | H5N1 | 2021-12-12 | AIV07-B1 |
| EPI_ISL_13370573 | A/chicken/England/070277/2021 | H5N1 | 2021-12-13 | AIV09 |
| EPI_ISL_13370603 | A/chicken/England/070359/2021 | H5N1 | 2021-12-13 | AIV09 |
| EPI_ISL_13370604 | A/chicken/England/070810/2021 | H5N1 | 2021-12-13 | AIV09 |
| EPI_ISL_13370605 | A/chicken/England/068020/2021 | H5N1 | 2021-12-14 | AIV07-B2 |
| EPI_ISL_13370606 | A/chicken/England/071013/2021 | H5N1 | 2021-12-14 | AIV07-B2 |
| EPI_ISL_13370607 | A/chicken/Scotland/070931/2021 | H5N1 | 2021-12-14 | AIV07-B2 |
| EPI_ISL_13370608 | A/chicken/England/072084/2021 | H5N1 | 2021-12-15 | AIV09 |
| EPI_ISL_13370609 | A/chicken/England/071631/2021 | H5N1 | 2021-12-15 | AIV09 |
| EPI_ISL_13370610 | A/chicken/England/072132/2021 | H5N1 | 2021-12-15 | AIV09 |
| EPI_ISL_13370611 | A/mute swan/Scotland/073112/2021 | H5N1 | 2021-12-15 | AIV07-B1 |
| EPI_ISL_13370612 | A/chicken/England/072378/2021 | H5N1 | 2021-12-16 | AIV09 |
| EPI_ISL_13370613 | A/chicken/England/073008/2021 | H5N1 | 2021-12-17 | AIV09 |
| EPI_ISL_13370614 | A/chicken/England/072926/2021 | H5N1 | 2021-12-17 | AIV07-B2 |
| EPI_ISL_13370615 | A/common buzzard/England/245061/2021 | H5N1 | 2021-12-17 | AIV07-B2 |
| EPI_ISL_13370616 | A/domestic duck/England/074574/2021 | H5N1 | 2021-12-21 | AIV07-B2 |

**Table S1 continued.**

| Accession Number | Isolate Name | Subtype | Collection Date | Genotype |
|---|---|---|---|---|
| EPI_ISL_13370617 | A/common buzzard/England/075831/2021 | H5N1 | 2021-12-21 | AIV07-B1 |
| EPI_ISL_8809153 | A/Muscovy duck/England/074477/2021 | H5N1 | 2021-12-21 | AIV07-B1 |
| EPI_ISL_13370618 | A/sparrowhawk/England/245233/2021 | H5N1 | 2021-12-22 | AIV07-B2 |
| EPI_ISL_13370619 | A/mute swan/England/387121/2021 | H5N1 | 2021-12-23 | AIV07-B2 |
| EPI_ISL_13370893 | A/Little gull/England/109275/2021 | H5N1 | 2021-12-24 | AIV09 |
| EPI_ISL_13370620 | A/turkey/England/075922/2021 | H5N1 | 2021-12-27 | AIV07-B2 |
| EPI_ISL_13370621 | A/turkey/England/075992/2021 | H5N1 | 2021-12-27 | AIV09 |
| EPI_ISL_13370622 | A/domestic duck/England/076200/2021 | H5N1 | 2021-12-29 | AIV07-B1 |
| EPI_ISL_13370623 | A/chicken/England/076198/2021 | H5N1 | 2021-12-29 | AIV07-B1 |
| EPI_ISL_13370624 | A/Whooper swan/England/000556/2021 | H5N1 | 2021-12-29 | AIV07-B1 |
| EPI_ISL_13370625 | A/domestic duck/England/076401/2021 | H5N1 | 2021-12-30 | AIV09 |
| EPI_ISL_13370626 | A/turkey/England/076717/2021 | H5N1 | 2021-12-30 | AIV09 |
| EPI_ISL_13370627 | A/mute swan/England/000030/2021 | H5N1 | 2021-12-31 | AIV07-B1 |
| EPI_ISL_13370628 | A/mute swan/England/244639/2021 | H5N1 | 2021-12-31 | AIV09 |
| EPI_ISL_13370629 | A/Common Buzzard/England/245285/2021 | H5N1 | 2021-12-31 | AIV07-B2 |
| EPI_ISL_13370630 | A/Goshawk/England/387180/2021 | H5N1 | 2021-12-31 | AIV07-B2 |
| EPI_ISL_13370631 | A/domestic duck/England/000357/2022 | H5N1 | 2022-01-02 | AIV09 |
| EPI_ISL_13370632 | A/chicken/England/000095/2022 | H5N1 | 2022-01-02 | AIV09 |
| EPI_ISL_11406401 | A/chicken/England/000187/2022 | H5N1 | 2022-01-03 | AIV07-B2 |
| EPI_ISL_13370633 | A/turkey/England/000733/2022 | H5N1 | 2022-01-04 | AIV09 |
| EPI_ISL_11406398 | A/chicken/England/002070/2022 | H5N1 | 2022-01-06 | AIV07-B2 |
| EPI_ISL_13370634 | A/chicken/England/002376/2022 | H5N1 | 2022-01-08 | AIV09 |
| EPI_ISL_13370679 | A/common gull/England/245481/2022 | H5N1 | 2022-01-08 | AIV09 |
| EPI_ISL_11406399 | A/turkey/England/004737/2022 | H5N1 | 2022-01-12 | AIV07-B2 |
| EPI_ISL_13370692 | A/turkey/England/004667/2022 | H5N1 | 2022-01-12 | AIV07-B2 |
| EPI_ISL_13370693 | A/Greylag goose/Sligoman/010298/2022 | H5N1 | 2022-01-14 | AIV07-B2 |
| EPI_ISL_13370694 | A/Domestic goose/England/006847/2022 | H5N1 | 2022-01-19 | AIV09 |
| EPI_ISL_13370695 | A/turkey/England/006963/2022 | H5N1 | 2022-01-20 | AIV07-B2 |
| EPI_ISL_11406402 | A/domestic duck/England/007588/2022 | H5N1 | 2022-01-23 | AIV07-B2 |
| EPI_ISL_13370696 | A/turkey/England/007586/2022 | H5N1 | 2022-01-23 | AIV07-B2 |
| EPI_ISL_13370697 | A/chicken/England/007584/2022 | H5N1 | 2022-01-23 | AIV07-B2 |
| EPI_ISL_13370698 | A/domestic goose/England/008781/2022 | H5N1 | 2022-01-26 | AIV07-B2 |
| EPI_ISL_13370699 | A/chicken/England/008775/2022 | H5N1 | 2022-01-26 | AIV07-B2 |
| EPI_ISL_13370700 | A/turkey/England/009687/2022 | H5N1 | 2022-01-27 | AIV07-B2 |
| EPI_ISL_13370702 | A/mallard duck/England/388009/2022 | H5N1 | 2022-01-27 | AIV07-B2 |
| EPI_ISL_13370701 | A/chicken/England/009811/2022 | H5N1 | 2022-01-27 | AIV07-B2 |
| EPI_ISL_13370894 | A/coot/England/388108/2022 | H5N1 | 2022-01-28 | AIV07-B2 |
| EPI_ISL_11406403 | A/black-headed gull/England/306270/2022 | H5N1 | 2022-01-31 | AIV07-B2 |
| EPI_ISL_11561589 | A/chicken/England/011981/2022 | H5N1 | 2022-02-02 | AIV07-B1 |
| EPI_ISL_11561593 | A/black-headed gull/England/388256/2022 | H5N1 | 2022-02-03 | AIV07-B2 |
| EPI_ISL_13370703 | A/domestic duck/England/012247/2022 | H5N1 | 2022-02-03 | AIV09 |
| EPI_ISL_13370895 | A/magpie/Scotland/012805/2022 | H5N1 | 2022-02-05 | AIV07-B2 |
| EPI_ISL_11561590 | A/chicken/England/012967/2022 | H5N1 | 2022-02-08 | AIV07-B2 |
| EPI_ISL_11561591 | A/domestic duck/England/012973/2022 | H5N1 | 2022-02-08 | AIV07-B2 |
| EPI_ISL_13370704 | A/chicken/England/014330/2022 | H5N1 | 2022-02-12 | AIV09 |
| EPI_ISL_11561592 | A/turkey/England/016515/2022 | H5N1 | 2022-02-20 | AIV20 |
| EPI_ISL_13370705 | A/pheasant/Wales/016441/2022 | H5N1 | 2022-02-20 | AIV09 |
| EPI_ISL_13370706 | A/pheasant/Wales/016303/2022 | H5N1 | 2022-02-20 | AIV09 |
| EPI_ISL_13370707 | A/domestic duck/England/017166/2022 | H5N1 | 2022-02-21 | AIV07-B2 |
| EPI_ISL_13370896 | A/Red-breasted goose/Jersey/018413/2022 | H5N1 | 2022-02-22 | AIV07-B2 |
| EPI_ISL_13370897 | A/Red-breasted goose/Jersey/018414/2022 | H5N1 | 2022-02-22 | AIV07-B2 |
| EPI_ISL_13370708 | A/domestic goose/England/019349/2022 | H5N1 | 2022-02-25 | AIV09 |
| EPI_ISL_13370709 | A/pheasant/England/019746/2022 | H5N1 | 2022-02-26 | AIV09 |
| EPI_ISL_13370710 | A/domestic duck/England/019894/2022 | H5N1 | 2022-02-28 | AIV09 |

### Table S1 continued.

| Accession Number | Isolate Name | Subtype | Collection Date | Genotype |
|---|---|---|---|---|
| EPI_ISL_13370907 | A/mute swan/Scotland/029918/2022 | H5N1 | 2022-03-07 | AIV09 |
| EPI_ISL_13370908 | A/mute swan/Scotland/029919/2022 | H5N1 | 2022-03-07 | AIV09 |
| EPI_ISL_13370909 | A/chicken/Scotland/024309/2022 | H5N1 | 2022-03-09 | AIV09 |
| EPI_ISL_13370910 | A/domestic duck/Scotland/024305/2022 | H5N1 | 2022-03-09 | AIV09 |
| EPI_ISL_13370911 | A/domestic duck/England/025226/2022 | H5N1 | 2022-03-10 | AIV09 |
| EPI_ISL_13370912 | A/domestic duck/England/025639/2022 | H5N1 | 2022-03-11 | AIV09 |
| EPI_ISL_13370913 | A/chicken/Scotland/029509/2022 | H5N1 | 2022-03-18 | AIV09 |
| EPI_ISL_13370914 | A/domestic duck/England/029617/2022 | H5N1 | 2022-03-20 | AIV09 |
| EPI_ISL_13370915 | A/domestic duck/England/032919/2022 | H5N1 | 2022-03-26 | AIV09 |
| EPI_ISL_13370916 | A/chicken/England/033318/2022 | H5N1 | 2022-03-27 | AIV09 |
| EPI_ISL_13370917 | A/chicken/England/034820/2022 | H5N1 | 2022-03-28 | AIV09 |
| EPI_ISL_13370918 | A/domestic duck/England/041295/2022 | H5N1 | 2022-04-04 | AIV09 |
| EPI_ISL_13370919 | A/domestic goose/England/041268/2022 | H5N1 | 2022-04-04 | AIV09 |
| EPI_ISL_13370920 | A/domestic duck/England/040831/2022 | H5N1 | 2022-04-04 | AIV07-B2 |
| EPI_ISL_13370921 | A/domestic duck/England/042266/2022 | H5N1 | 2022-04-06 | AIV09 |
| EPI_ISL_13370922 | A/chicken/England/042391/2022 | H5N1 | 2022-04-06 | AIV07-B2 |
| EPI_ISL_13370923 | A/chicken/England/046274/2022 | H5N1 | 2022-04-12 | AIV09 |
| EPI_ISL_13370924 | A/chicken/England/053826/2022 | H5N1 | 2022-04-21 | AIV07-B2 |
| EPI_ISL_13370925 | A/chicken/England/063896/2022 | H5N1 | 2022-05-05 | AIV07-B2 |

**Table S2.** All polymorphisms associated with altered virulence, host susceptibility and antiviral resistance for the different H5 AIV subtypes detected in the UK between 2020-2022. All different polymorphisms observed for each subtype are shown, along with their previously demonstrated phenotype.

| Protein | Amino Acid Substitution | Phenotype | Reference | Sequences containing the polymorphism (%) | | | | | |
|---|---|---|---|---|---|---|---|---|---|
| | | | | H5N1 (N=200) | H5N2 (N=1) | H5N3 LPAIV (N=1) | H5N3 HPAIV (N=1) | H5N5 (N=3) | H5N8 (N=33) |
| HA (H5 numbering) | S107R, T108I | Increased virulence in chickens and mice, increased pH of fusion | PMID: 31428925, 29899102 | 100 | 0 | 0 | 100 | 100 | 100 |
| | S123P, R497K | Increased virus binding to α2-6 | PMID: 31428925, 17108965 | 1 | 0 | 0 | 0 | 0 | 0 |
| | S133A | Increased pseudovirus binding to α2-6 | PMID: 31428925, 17690300 | 100 | 0 | 0 | 100 | 100 | 100 |
| | S154N | Increased virus binding to α2-6 | PMID: 31428925, 20427525 | 100 | 100 | 0 | 100 | 100 | 100 |
| | S155N | Increased virus binding to α2-6 | PMID: 31428925, 20427525 | 2.5 | 100 | 100 | 0 | 0 | 0 |
| | T156A | Increased virus binding to α2-6 and increased transmission in guinea pigs | PMID: 31428925, 20427525, 20041223 | 100 | 100 | 100 | 100 | 100 | 100 |
| | V182N | Increased binding to α2–6, decreased binding to α2–3 | PMID: 31428925, 23760233 | 100 | 100 | 100 | 100 | 100 | 100 |
| | K218Q, S223R | Increased virus binding to α2–3 and α2–6 | PMID: 31428925, 27869615 | 100 | 0 | 0 | 100 | 100 | 100 |
| | E251K | Increased virus binding to α2-6 | PMID: 31428925, 22056389 | 0.5 | 0 | 0 | 0 | 0 | 0 |
| | K394E | Increased pH of fusion, decreased HA stability, decreased virulence in mice | PMID: 31428925, 31325838 | 100 | 0 | 100 | 100 | 100 | 100 |
| NA (N2 Numbering) | I117T | Reduced susceptibility to oseltamivir and zanamivir | PMID: 31428925, 31282375 | 0 | 100 | 100 | 100 | 0 | 0 |
| | A401T | Increased virus binding to α2-3 | PMID: 31428925, 28202753 | 100 | 0 | 0 | 0 | 0 | 0 |
| PB2 | T63I [with PB1: M677T] | Pathogenic in mice | PMID: 21371335 | 100 | 100 | 100 | 100 | 100 | 100 |
| | L89V, G309D, T339K, R477G, I495V, K627E, A676T | Increased polymerase activity in mammalian cell line and increased virulence in mice | PMID: 31428925, 19393699 | 96.5 | 100 | 100 | 100 | 100 | 96.97 |
| | E249G | Increased polymerase activity | PMID: 29593225 | 1.5 | 0 | 0 | 0 | 0 | 0 |
| | K251R | Increased virulence in mice | PMID: 26829383 | 99 | 100 | 100 | 100 | 100 | 100 |
| | I292V | Increased polymerase activity in mammalian cell line, increased virulence in mice | PMID: 31428925, 31305236, 26782141 | 0 | 0 | 100 | 100 | 100 | 100 |
| | G309D | Enhanced polymerase activity, increased virulence in mice | PMID: 32709116, 19393699 | 100 | 100 | 100 | 100 | 100 | 100 |
| | T339K | Enhanced polymerase activity, increased virulence in mice | PMID: 32709116, 19393699 | 100 | 100 | 100 | 100 | 100 | 96.97 |
| | Q368R | Increased polymerase activity, increased virulence in mammals | PMID: 32709116, 16533883 | 99.5 | 100 | 100 | 100 | 100 | 100 |
| | K389R | Increased polymerase activity and replication in mammalian cell line | PMID: 31428925, 27889648 | 99.5 | 100 | 100 | 100 | 0 | 100 |

**Table S2 continued.**

| Protein | Amino Acid Substitution | Phenotype | Reference | Sequences containing the polymorphism (%) | | | | | |
|---|---|---|---|---|---|---|---|---|---|
| | | | | H5N1 (N=200) | H5N2 (N=1) | H5N3 LPAIV (N=1) | H5N3 HPAIV (N=1) | H5N5 (N=3) | H5N8 (N=33) |
| **PB2** | H447Q | Increased polymerase activity, increased virulence in mammals | PMID: 32709116, 16533883 | 100 | 100 | 100 | 100 | 100 | 100 |
| | R477G | Enhanced polymerase activity, increased virulence in mice | PMID: 32709116, 19393699 | 100 | 100 | 100 | 100 | 100 | 100 |
| | I495V | Enhanced polymerase activity, increased virulence in mice | PMID: 32709116, 19393699 | 100 | 100 | 100 | 100 | 100 | 100 |
| | V598T | Increased polymerase activity and replication in mammalian cells, increased virulence in mice | PMID: 31428925, 27889648 | 100 | 100 | 100 | 100 | 100 | 100 |
| | K627E | Increased virulence in chickens | PMID: 31428925, 22363523 | 99.5 | 100 | 100 | 100 | 100 | 100 |
| | E627K | Increased polymerase activity and replication in mammalian cell line, increase virulence in mice; contributes to airborne pathogenicity of IAVs in ferrets and contact transmission in guinea pigs; decreases polymerase activity and replication in avian cell lines; decreases virulence in chickens | PMID: 31428925, 31081750, 22723413, 17922570, 11546875, 21849466, 23843645, 28775271, 20016035, 16228009, 19052090, 17521765, 15016548, 19692471, 17098982, 21846828, 25782865, 26560088, 25194918, 24007444, 24394699 | 0.5 | 0 | 0 | 0 | 0 | 0 |
| | A676T | Enhanced polymerase active, increased virulence in mice | PMID: 32709116, 19393699 | 99 | 100 | 100 | 100 | 100 | 100 |
| | D701N | Increased polymerase activity, increased virulence in mammals; enhanced replication efficiency, increased virulence and contact transmission in guinea pigs, increased virulence in mice | PMID: 31428925, 31111259, 19119420, 26082035, 20041223, 19264775, 16140781, 26845764, 24403592, 26560088 | 0 | 0 | 0 | 0 | 0 | 6.06 |
| **PB1** | D3V | Increased polymerase activity and viral replication in avian and mammalian cell lines | PMID: 31428925, 27926816 | 97 | 100 | 100 | 100 | 33.33 | 96.97 |
| | D622G | Increased polymerase activity and virulence in mice | PMID: 31428925, 26656683 | 100 | 100 | 100 | 100 | 100 | 100 |
| **PB1-F2** | T51M | Decrease polymerase activity, replication and virulence in ducks | PMID: 20383540 | 0.5 | 0 | 0 | 0 | 0 | 0 |
| | N66S | Enhanced replication, virulence and antiviral response in mice | PMID: 31428925, 21852950, 17922571 | 100 | 100 | 100 | 100 | 0 | 0 |
| | Truncated PB1-F2 | Increased polymerase activity and virulence in mice | PMID: 25787281 | 0 | 0 | 0 | 0 | 100 | 100 |
| **PA** | S37A | Increased polymerase activity in mammalian cell line | PMID: 31428925, 24371069 | 100 | 100 | 100 | 100 | 100 | 100 |
| | V63I | Increase polymerase activity and enhanced replication in mammalian cell line, increased virulence in mice | PMID: 31428925, 28113045, 27886255 | 0.5 | 0 | 0 | 0 | 0 | 0 |

**Table S2 continued.**

| Protein | Amino Acid Substitution | Phenotype | Reference | Sequences containing the polymorphism (%) | | | | | |
|---|---|---|---|---|---|---|---|---|---|
| | | | | H5N1 (N=200) | H5N2 (N=1) | H5N3 LPAIV (N=1) | H5N3 HPAIV (N=1) | H5N5 (N=3) | H5N8 (N=33) |
| **PA** | K142N | Increased virulence in mice | PMID: 31428925, 20016035 | 0.5 | 0 | 0 | 0 | 0 | 0 |
| | P190S | Decreased virulence in mice | PMID: 31428925, 27105450 | 100 | 100 | 100 | 100 | 100 | 100 |
| | P224P, N383D | Increased polymerase activity and enhanced viral replication in duck and mouse cell lines, increased virulence in mice and ducks | PMID: 31428925, 26000865, 21177821 | 2.5 | 0 | 0 | 0 | 0 | 0 |
| | N383D | Increased polymerase activity in mammalian and avian cell lines | PMID: 31428925, 26000865, 21177821 | 100 | 100 | 100 | 100 | 100 | 100 |
| | Q400P | Decreased virulence in mice | PMID: 31428925, 27105450 | 43 | 0 | 100 | 0 | 0 | 0 |
| | N409S | Increased polymerase activity and replication in mammalian cell line | PMID: 31428925, 24371069 | 100 | 100 | 100 | 100 | 100 | 100 |
| **NP** | I41V | Increased polymerase activity in mammalian cell line | PMID: 31428925, 25940512 | 0.5 | 0 | 0 | 100 | 0 | 0 |
| | M105V | Increased virulence in chickens | PMID: 31428925, 21123376, 21795332 | 57.5 | 100 | 0 | 0 | 100 | 100 |
| | A184K | Increased replication in avian cells and virulence in chickens, enhanced IFN response | PMID: 31428925, 19475480 | 100 | 100 | 100 | 100 | 100 | 100 |
| | N319K | Increased polymerase activity and replication in mammalian cell line | PMID: 31428925, 16339318, 18248089 | 5.5 | 0 | 0 | 0 | 0 | 0 |
| **M1** | N30D | Increased virulence in mice | PMID: 31428925, 19117585 | 100 | 100 | 100 | 100 | 100 | 100 |
| | I43M | Increased virulence in mice, chickens and ducks | PMID: 31428925, 26368015 | 100 | 100 | 100 | 100 | 100 | 100 |
| | T139A | Increased virulence in mice | PMID: 8879138, 10426210 | 0.5 | 0 | 0 | 0 | 0 | 0 |
| | T215A | Increased virulence in mice | PMID: 31428925, 19117585 | 0 | 100 | 100 | 100 | 100 | 100 |
| **M2** | A30S | Reduced susceptibility to amantadine and rimantadine | PMID: 31428925, 15673732, 20834097, 16703504, 2723453, 16081121, | 0.5 | 0 | 0 | 0 | 0 | 0 |
| **NS1** | P42S | Increased virulence in mice | PMID: 31428925, 18032512 | 100 | 0 | 100 | 100 | 100 | 100 |
| | L103F, I106M | Increased virulence in mice | PMID: 31428925, 19052083, 21593152 | 100 | 0 | 100 | 100 | 100 | 90.91 |
| | I106M | Increased viral replication in mammalian cells virulence in mice | PMID: 31428925, 25078692 | 100 | 100 | 100 | 100 | 100 | 93.94 |
| | C138F | Increased replication in mammalian cells, decreased interferon response | PMID: 31428925, 29677653 | 99 | 100 | 100 | 100 | 100 | 100 |
| | V149A | Increased virulence and decreased interferon response in chickens | PMID: 31428925, 16971424 | 100 | 100 | 100 | 100 | 100 | 100 |
| | 222-230 Deletion | Increased replication in mammalian and avian cell lines | PMID: 31428925, 20686040 | 0 | 0 | 0 | 0 | 100 | 100 |

**Table S2 continued.**

| Protein | Amino Acid Substitution | Phenotype | Reference | Sequences containing the polymorphism (%) | | | | | |
|---|---|---|---|---|---|---|---|---|---|
| | | | | H5N1 (N=200) | H5N2 (N=1) | H5N3 LPAIV (N=1) | H5N3 HPAIV (N=1) | H5N5 (N=3) | H5N8 (N=33) |
| **NS1** | 227-ESEV-230 (PDZ Domain) | Increased virulence in mice; Decreased viral replication in mammalian and avian cell lines; decreased viral replication in human and duck cell lines | PMID: 31428925, 18334632, 20410267, 20686040 | 100 | 100 | 100 | 100 | 100 | 100 |

**Table S3.** Sequences used to investigate lateral spread between infected premises (IP) and wild birds (WB) in the different geographic clusters. The collection date and associated H5 subtype/genotype, as well as the reference sequence used for each cluster are also provided.

| Infected Premises/Wild Bird Number | Sequence | Collection Date | H5Nx Subtype/Genotype |
|---|---|---|---|
| **Cluster 1** | | | |
| IP1 | A/turkey/England/037784/2020 | 2020-11-28 | H5N8 |
| IP2 | A/turkey/England/038115/2020 | 2020-12-02 | H5N8 |
| Reference | A/Greylag goose/England/032698/2020 | 2020-11-03 | H5N8 |
| **Cluster 2** | | | |
| IP1 | A/chicken/England/057016/2021 | 2021-11-12 | H5N1 AIV09 |
| IP2 | A/turkey/England/059498/2021 | 2021-11-20 | H5N1 AIV09 |
| IP3 | A/chicken/England/59973/2021 | 2021-11-20 | H5N1 AIV09 |
| IP4 | A/turkey/England/059870/2021 | 2021-11-21 | H5N1 AIV09 |
| IP5 | A/chicken/England/062137/2021 | 2021-11-25 | H5N1 AIV09 |
| IP6 | A/turkey/England/062369/2021 | 2021-11-27 | H5N1 AIV09 |
| IP7 | A/chicken/England/064357/2021 | 2021-12-01 | H5N1 AIV09 |
| IP8 | A/chicken/England/065487/2021 | 2021-12-04 | H5N1 AIV09 |
| IP9 | A/turkey/England/067429/2021 | 2021-12-08 | H5N1 AIV09 |
| Reference | A/chicken/Scotland/054477/2021 | 2021-11-01 | H5N1 AIV09 |
| **Cluster 3** | | | |
| WB1 | A/mute swan/England/243983/2021 | 2021-11-13 | H5N1 AIV09 |
| IP1 | A/chicken/England/061210/2021 | 2021-11-22 | H5N1 AIV09 |
| IP2 | A/chicken/England/063664/2021 | 2021-11-28 | H5N1 AIV09 |
| IP3 | A/chicken/England/066019/2021 | 2021-12-04 | H5N1 AIV09 |
| IP4 | A/chicken/England/066782/2021 | 2021-12-05 | H5N1 AIV09 |
| IP5 | A/chicken/England/070810/2021 | 2021-12-13 | H5N1 AIV09 |
| IP6 | A/turkey/England/076717/2021 | 2021-12-30 | H5N1 AIV09 |
| WB2 | A/common gull/England/245481/2022 | 2022-01-08 | H5N1 AIV09 |
| Reference | A/chicken/Scotland/054477/2021 | 2021-11-01 | H5N1 AIV09 |
| **Cluster 4** | | | |
| IP1 | A/chicken/England/069549/2021 | 2021-12-11 | H5N1 AIV09 |
| IP2 | A/chicken/England/070105/2021 | 2021-12-11 | H5N1 AIV09 |
| IP3 | A/chicken/England/070277/2021 | 2021-12-13 | H5N1 AIV09 |
| IP4 | A/chicken/England/070359/2021 | 2021-12-13 | H5N1 AIV09 |
| IP5 | A/chicken/England/072084/2021 | 2021-12-15 | H5N1 AIV09 |
| IP6 | A/chicken/England/071631/2021 | 2021-12-15 | H5N1 AIV09 |
| IP7 | A/chicken/England/072132/2021 | 2021-12-15 | H5N1 AIV09 |
| IP8 | A/chicken/England/072378/2021 | 2021-12-16 | H5N1 AIV09 |
| IP9 | A/chicken/England/073008/2021 | 2021-12-17 | H5N1 AIV09 |
| IP10 | A/turkey/England/075992/2021 | 2021-12-27 | H5N1 AIV09 |
| IP11 | A/domestic duck/England/076401/2021 | 2021-12-30 | H5N1 AIV09 |
| IP12 | A/domestic duck/England/000357/2022 | 2022-01-02 | H5N1 AIV09 |
| IP12 | A/chicken/England/000095/2022 | 2022-01-02 | H5N1 AIV09 |
| IP13 | A/turkey/England/000733/2022 | 2022-01-04 | H5N1 AIV09 |
| IP14 | A/chicken/England/002376/2022 | 2022-01-08 | H5N1 AIV09 |
| Reference | A/chicken/Scotland/054477/2021 | 2021-11-01 | H5N1 AIV09 |
| **Cluster 5** | | | |
| IP1 | A/chicken/England/072926/2021 | 2021-12-17 | H5N1 AIV07-B2 |
| IP2 | A/domestic duck/England/074574/2021 | 2021-12-21 | H5N1 AIV07-B2 |
| WB1 | A/mute swan/England/387121/2021 | 2021-12-23 | H5N1 AIV07-B2 |
| IP3 | A/turkey/England/004737/2022 | 2022-01-12 | H5N1 AIV07-B2 |
| IP4 | A/turkey/England/004667/2022 | 2022-01-12 | H5N1 AIV07-B2 |
| IP5 | A/turkey/England/006963/2022 | 2022-01-20 | H5N1 AIV07-B2 |
| IP6 | A/turkey/England/009687/2022 | 2022-01-27 | H5N1 AIV07-B2 |

**Table S3 continued.**

| Infected Premises/Wild Bird Number | Sequence | Collection Date | H5Nx Subtype/Genotype |
|---|---|---|---|
| Reference | A/Greylag goose/England/054503/2021 | 2021-10-30 | H5N1 AIV07-B2 |
| **Cluster 6** | | | |
| IP1 | A/domestic goose/England/019349/2022 | 2022-02-25 | H5N1 AIV09 |
| IP2 | A/domestic duck/England/019894/2022 | 2022-02-28 | H5N1 AIV09 |
| IP3 | A/domestic duck/England/025226/2022 | 2022-03-10 | H5N1 AIV09 |
| IP4 | A/domestic duck/England/025639/2022 | 2022-03-11 | H5N1 AIV09 |
| IP5 | A/domestic duck/England/029617/2022 | 2022-03-20 | H5N1 AIV09 |
| IP6 | A/domestic duck/England/032919/2022 | 2022-03-26 | H5N1 AIV09 |
| IP7 | A/chicken/England/033318/2022 | 2022-03-27 | H5N1 AIV09 |
| IP8 | A/chicken/England/034820/2022 | 2022-03-28 | H5N1 AIV09 |
| Reference | A/chicken/Scotland/054477/2021 | 2021-11-01 | H5N1 AIV09 |
| **Cluster 7** | | | |
| IP1 | A/domestic duck/England/041295/2022 | 2022-04-04 | H5N1 AIV09 |
| IP1 | A/domestic goose/England/041268/2022 | 2022-04-04 | H5N1 AIV09 |
| IP2 | A/domestic duck/England/042266/2022 | 2022-04-06 | H5N1 AIV09 |
| IP3 | A/chicken/England/046274/2022 | 2022-04-12 | H5N1 AIV09 |
| Reference | A/chicken/Scotland/054477/2021 | 2021-11-01 | H5N1 AIV09 |

**Table S4**. Outputs of the BSSVS analysis for Cluster 1 showing the rates of transmission between sampled infected premises (IP).

| Source | Target | Bayes Factor | Posterior Probability |
|---|---|---|---|
| IP1 | IP2 | 1.5237 | 0.9304 |
| IP1 | H5N8 Reference | 0.4715 | 0.8054 |
| IP2 | H5N8 Reference | 0.3894 | 0.7736 |

**Table S5.** Outputs of the BSSVS analysis for Cluster 2 showing the rates of transmission between sampled infected premises (IP).

| Source | Target | Bayes Factor | Posterior Probability |
|---|---|---|---|
| AIV09 Reference | IP1 | 2.2777 | 0.3847 |
| AIV09 Reference | IP2 | 0.8395 | 0.1873 |
| AIV09 Reference | IP3 | 1.3459 | 0.2698 |
| AIV09 Reference | IP4 | 0.8118 | 0.1823 |
| AIV09 Reference | IP5 | 3.6142 | 0.4981 |
| AIV09 Reference | IP6 | 0.5831 | 0.1380 |
| AIV09 Reference | IP7 | 0.4450 | 0.1089 |
| AIV09 Reference | IP8 | 0.4356 | 0.1068 |
| AIV09 Reference | IP9 | 0.4275 | 0.1050 |
| IP1 | IP2 | 1.8447 | 0.3362 |
| IP1 | IP3 | 4.6024 | 0.5582 |
| IP1 | IP4 | 1.7718 | 0.3272 |
| IP1 | IP5 | 1.3649 | 0.2726 |
| IP1 | IP6 | 0.8960 | 0.1974 |
| IP1 | IP7 | 0.6324 | 0.1479 |
| IP1 | IP8 | 0.5817 | 0.1377 |
| IP1 | IP9 | 0.5828 | 0.1379 |
| IP2 | IP3 | 1.1240 | 0.2358 |
| IP2 | IP4 | 2.4172 | 0.3989 |
| IP2 | IP5 | 0.6363 | 0.1487 |
| IP2 | IP6 | 1.0631 | 0.2259 |
| IP2 | IP7 | 3.8497 | 0.5138 |
| IP2 | IP8 | 2.8248 | 0.4368 |
| IP2 | IP9 | 2.8140 | 0.4358 |
| IP3 | IP4 | 1.1323 | 0.2371 |
| IP3 | IP5 | 0.9382 | 0.2048 |
| IP3 | IP6 | 0.6857 | 0.1584 |
| IP3 | IP7 | 0.5111 | 0.1230 |
| IP3 | IP8 | 0.4818 | 0.1168 |
| IP3 | IP9 | 0.4735 | 0.1150 |
| IP4 | IP5 | 0.6285 | 0.1472 |
| IP4 | IP6 | 11.8116 | 0.7643 |
| IP4 | IP7 | 0.7462 | 0.1700 |
| IP4 | IP8 | 0.6497 | 0.1514 |
| IP4 | IP9 | 0.6196 | 0.1454 |
| IP5 | IP6 | 0.5019 | 0.1211 |
| IP5 | IP7 | 0.4134 | 0.1019 |
| IP5 | IP8 | 0.4019 | 0.0994 |
| IP5 | IP9 | 0.4012 | 0.0992 |
| IP6 | IP7 | 0.5000 | 0.1207 |
| IP6 | IP8 | 0.4532 | 0.1107 |
| IP6 | IP9 | 0.4629 | 0.1128 |
| IP7 | IP8 | 3.8190 | 0.5118 |
| IP7 | IP9 | 2.3083 | 0.3879 |
| IP8 | IP9 | 1.6855 | 0.3164 |

**Table S6.** Outputs of the BSSVS analysis for Cluster 3 showing the rates of transmission between sampled infected premises (IP) and/or wild birds (WB).

| Source | Target | Bayes Factor | Posterior Probability |
|---|---|---|---|
| AIV09 Reference | IP1 | 0.9810 | 0.2380 |
| AIV09 Reference | IP2 | 0.6946 | 0.1811 |
| AIV09 Reference | IP3 | 0.6476 | 0.1709 |
| AIV09 Reference | IP4 | 0.5916 | 0.1585 |
| AIV09 Reference | IP5 | 0.9948 | 0.2405 |
| AIV09 Reference | IP6 | 1.6239 | 0.3408 |
| AIV09 Reference | WB1 | 2.8156 | 0.4727 |
| AIV09 Reference | WB2 | 0.9005 | 0.2228 |
| IP1 | IP2 | 4.5138 | 0.5897 |
| IP1 | IP3 | 5.2849 | 0.6272 |
| IP1 | IP4 | 2.5283 | 0.4460 |
| IP1 | IP5 | 1.6775 | 0.3481 |
| IP1 | IP6 | 0.6975 | 0.1817 |
| IP1 | WB1 | 1.0058 | 0.2425 |
| IP1 | WB2 | 1.2356 | 0.2823 |
| IP2 | IP3 | 1.4667 | 0.3183 |
| IP2 | IP4 | 4.2934 | 0.5775 |
| IP2 | IP5 | 0.9747 | 0.2368 |
| IP2 | IP6 | 0.5618 | 0.1517 |
| IP2 | WB1 | 0.6806 | 0.1781 |
| IP2 | WB2 | 0.8486 | 0.2127 |
| IP3 | IP4 | 1.0731 | 0.2546 |
| IP3 | IP5 | 0.8994 | 0.2226 |
| IP3 | IP6 | 0.5310 | 0.1446 |
| IP3 | WB1 | 0.6413 | 0.1695 |
| IP3 | WB2 | 0.7920 | 0.2014 |
| IP4 | IP5 | 0.7863 | 0.2002 |
| IP4 | IP6 | 0.5040 | 0.1383 |
| IP4 | WB1 | 0.5769 | 0.1551 |
| IP4 | WB2 | 0.7261 | 0.1878 |
| IP5 | IP6 | 0.7514 | 0.1930 |
| IP5 | WB1 | 0.9909 | 0.2398 |
| IP5 | WB2 | 1.6657 | 0.3465 |
| IP6 | WB1 | 2.6604 | 0.4586 |
| IP6 | WB2 | 0.6606 | 0.1738 |
| WB1 | WB2 | 0.8310 | 0.2092 |

**Table S7.** Outputs of the BSSVS analysis for Cluster 4 showing the rates of transmission between sampled infected premises (IP).

| Source | Target | Bayes Factor | Posterior Probability |
|---|---|---|---|
| AIV09 Reference | IP1 | 1.3217 | 0.1770 |
| AIV09 Reference | IP2 | 0.8517 | 0.1217 |
| AIV09 Reference | IP3 | 0.7742 | 0.1119 |
| AIV09 Reference | IP4 | 1.0586 | 0.1469 |
| AIV09 Reference | IP5 | 0.6476 | 0.0953 |
| AIV09 Reference | IP6 | 7.1427 | 0.5375 |
| AIV09 Reference | IP7 | 0.7583 | 0.1098 |
| AIV09 Reference | IP8 | 0.6243 | 0.0922 |
| AIV09 Reference | IP9 | 0.7163 | 0.1044 |
| AIV09 Reference | IP10 | 0.4578 | 0.0693 |
| AIV09 Reference | IP11 | 0.3950 | 0.0604 |
| AIV09 Reference | IP12 | 0.3646 | 0.0560 |
| AIV09 Reference | IP13 | 0.6293 | 0.0929 |
| AIV09 Reference | IP14 | 0.3382 | 0.0522 |
| IP1 | IP2 | 2.3872 | 0.2797 |
| IP1 | IP3 | 1.7445 | 0.2211 |
| IP1 | IP4 | 2.4978 | 0.2890 |
| IP1 | IP5 | 1.3328 | 0.1782 |
| IP1 | IP6 | 0.9464 | 0.1334 |
| IP1 | IP7 | 1.7813 | 0.2247 |
| IP1 | IP8 | 1.0897 | 0.1506 |
| IP1 | IP9 | 1.6003 | 0.2066 |
| IP1 | IP10 | 0.7574 | 0.1097 |
| IP1 | IP11 | 0.5848 | 0.0869 |
| IP1 | IP12 | 0.5550 | 0.0828 |
| IP1 | IP13 | 1.0992 | 0.1517 |
| IP1 | IP14 | 0.4626 | 0.0700 |
| IP2 | IP3 | 1.7620 | 0.2228 |
| IP2 | IP4 | 2.1492 | 0.2591 |
| IP2 | IP5 | 1.3262 | 0.1775 |
| IP2 | IP6 | 0.6892 | 0.1008 |
| IP2 | IP7 | 2.1999 | 0.2636 |
| IP2 | IP8 | 15.4176 | 0.7150 |
| IP2 | IP9 | 1.8628 | 0.2326 |
| IP2 | IP10 | 0.8468 | 0.1211 |
| IP2 | IP11 | 0.6165 | 0.0912 |
| IP2 | IP12 | 0.6177 | 0.0913 |
| IP2 | IP13 | 1.1106 | 0.1530 |
| IP2 | IP14 | 0.4923 | 0.0742 |
| IP3 | IP4 | 1.9536 | 0.2412 |
| IP3 | IP5 | 10.8628 | 0.6387 |
| IP3 | IP6 | 0.6123 | 0.0906 |
| IP3 | IP7 | 1.6621 | 0.2129 |
| IP3 | IP8 | 0.8699 | 0.1240 |
| IP3 | IP9 | 1.3750 | 0.1828 |
| IP3 | IP10 | 0.7842 | 0.1132 |
| IP3 | IP11 | 0.6066 | 0.0898 |
| IP3 | IP12 | 0.5688 | 0.0847 |
| IP3 | IP13 | 0.9260 | 0.1309 |
| IP3 | IP14 | 0.4066 | 0.0620 |
| IP4 | IP5 | 1.5055 | 0.1968 |
| IP4 | IP6 | 0.8239 | 0.1182 |
| IP4 | IP7 | 1.7938 | 0.2259 |
| IP4 | IP8 | 0.9719 | 0.1365 |
| IP4 | IP9 | 1.5809 | 0.2046 |

**Table S7 continued.**

| Source | Target | Bayes Factor | Posterior Probability |
|---|---|---|---|
| IP4 | IP10 | 0.7742 | 0.1119 |
| IP4 | IP11 | 0.5926 | 0.0879 |
| IP4 | IP12 | 0.5971 | 0.0885 |
| IP4 | IP13 | 1.0703 | 0.1483 |
| IP4 | IP14 | 0.4563 | 0.0691 |
| IP5 | IP6 | 0.5575 | 0.0832 |
| IP5 | IP7 | 1.2384 | 0.1677 |
| IP5 | IP8 | 0.7095 | 0.1035 |
| IP5 | IP9 | 1.0845 | 0.1500 |
| IP5 | IP10 | 0.6752 | 0.0990 |
| IP5 | IP11 | 0.5227 | 0.0784 |
| IP5 | IP12 | 0.5063 | 0.0761 |
| IP5 | IP13 | 0.8819 | 0.1255 |
| IP5 | IP14 | 0.4085 | 0.0623 |
| IP6 | IP7 | 0.6074 | 0.0899 |
| IP6 | IP8 | 0.5445 | 0.0814 |
| IP6 | IP9 | 0.5709 | 0.0850 |
| IP6 | IP10 | 0.4132 | 0.0630 |
| IP6 | IP11 | 0.3405 | 0.0525 |
| IP6 | IP12 | 0.3428 | 0.0528 |
| IP6 | IP13 | 0.5304 | 0.0794 |
| IP6 | IP14 | 0.3193 | 0.0494 |
| IP7 | IP8 | 0.9582 | 0.1349 |
| IP7 | IP9 | 2.7911 | 0.3123 |
| IP7 | IP10 | 1.1407 | 0.1565 |
| IP7 | IP11 | 0.8530 | 0.1219 |
| IP7 | IP12 | 0.8336 | 0.1194 |
| IP7 | IP13 | 1.6798 | 0.2146 |
| IP7 | IP14 | 0.5881 | 0.0873 |
| IP8 | IP9 | 0.8797 | 0.1252 |
| IP8 | IP10 | 0.5027 | 0.0756 |
| IP8 | IP11 | 0.4027 | 0.0615 |
| IP8 | IP12 | 0.4136 | 0.0630 |
| IP8 | IP13 | 0.6892 | 0.1008 |
| IP8 | IP14 | 0.3443 | 0.0530 |
| IP9 | IP10 | 0.9953 | 0.1394 |
| IP9 | IP11 | 0.7522 | 0.1090 |
| IP9 | IP12 | 0.7069 | 0.1032 |
| IP9 | IP13 | 1.4410 | 0.1899 |
| IP9 | IP14 | 0.5538 | 0.0827 |
| IP10 | IP11 | 3.9242 | 0.3897 |
| IP10 | IP12 | 4.4988 | 0.4226 |
| IP10 | IP13 | 0.7742 | 0.1119 |
| IP10 | IP14 | 2.1213 | 0.2566 |
| IP11 | IP12 | 3.7717 | 0.3803 |
| IP11 | IP13 | 0.5836 | 0.0867 |
| IP11 | IP14 | 8.1655 | 0.5705 |
| IP12 | IP13 | 0.5530 | 0.0825 |
| IP12 | IP14 | 2.0997 | 0.2546 |
| IP13 | IP14 | 0.4515 | 0.0684 |

**Table S8.** Outputs of the BSSVS analysis for Cluster 5 showing the rates of transmission between sampled infected premises (IP) and/or wild birds (WB).

| Source | Target | Bayes Factor | Posterior Probability |
|---|---|---|---|
| AIV07-B2 Reference | IP1 | 0.9562 | 0.2659 |
| AIV07-B2 Reference | IP2 | 0.8564 | 0.2450 |
| AIV07-B2 Reference | IP3 | 1.6210 | 0.3805 |
| AIV07-B2 Reference | IP4 | 1.6240 | 0.3809 |
| AIV07-B2 Reference | IP5 | 2.1597 | 0.4500 |
| AIV07-B2 Reference | IP6 | 1.5563 | 0.3709 |
| AIV07-B2 Reference | WB1 | 2.1767 | 0.4519 |
| IP1 | IP2 | 24.5136 | 0.9028 |
| IP1 | IP3 | 0.5676 | 0.1770 |
| IP1 | IP4 | 0.5774 | 0.1795 |
| IP1 | IP5 | 0.7516 | 0.2216 |
| IP1 | IP6 | 0.5549 | 0.1737 |
| IP1 | WB1 | 0.6511 | 0.1979 |
| IP2 | IP3 | 0.5560 | 0.1740 |
| IP2 | IP4 | 0.5329 | 0.1680 |
| IP2 | IP5 | 0.6718 | 0.2029 |
| IP2 | IP6 | 0.5444 | 0.1710 |
| IP2 | WB1 | 0.6062 | 0.1868 |
| IP3 | IP4 | 4.8166 | 0.6460 |
| IP3 | IP5 | 0.9302 | 0.2606 |
| IP3 | IP6 | 0.7970 | 0.2319 |
| IP3 | WB1 | 0.9687 | 0.2685 |
| IP4 | IP5 | 0.9440 | 0.2634 |
| IP4 | IP6 | 0.7854 | 0.2293 |
| IP4 | WB1 | 0.9734 | 0.2694 |
| IP5 | IP6 | 0.9262 | 0.2597 |
| IP5 | WB1 | 1.1091 | 0.2959 |
| IP6 | WB1 | 3.6835 | 0.5825 |

**Table S9**. Outputs of the BSSVS analysis for Cluster 6 showing the rates of transmission between sampled infected premises (IP).

| **Source** | **Target** | **Bayes Factor** | **Posterior Probability** |
|---|---|---|---|
| AIV09 Reference | IP1 | 3.2527 | 0.5087 |
| AIV09 Reference | IP2 | 1.2594 | 0.2862 |
| AIV09 Reference | IP3 | 0.8030 | 0.2036 |
| AIV09 Reference | IP4 | 0.7716 | 0.1972 |
| AIV09 Reference | IP5 | 0.5101 | 0.1397 |
| AIV09 Reference | IP6 | 0.4530 | 0.1260 |
| AIV09 Reference | IP7 | 0.4059 | 0.1144 |
| AIV09 Reference | IP8 | 0.4260 | 0.1194 |
| IP1 | IP2 | 1.5129 | 0.3251 |
| IP1 | IP3 | 0.8085 | 0.2047 |
| IP1 | IP4 | 0.7597 | 0.1948 |
| IP1 | IP5 | 0.4876 | 0.1344 |
| IP1 | IP6 | 0.4425 | 0.1235 |
| IP1 | IP7 | 0.4095 | 0.1153 |
| IP1 | IP8 | 0.4139 | 0.1164 |
| IP2 | IP3 | 6.3674 | 0.6696 |
| IP2 | IP4 | 5.7921 | 0.6484 |
| IP2 | IP5 | 1.6853 | 0.3492 |
| IP2 | IP6 | 1.0167 | 0.2445 |
| IP2 | IP7 | 0.8568 | 0.2143 |
| IP2 | IP8 | 0.7978 | 0.2025 |
| IP3 | IP4 | 1.9804 | 0.3867 |
| IP3 | IP5 | 0.8650 | 0.2159 |
| IP3 | IP6 | 0.6069 | 0.1619 |
| IP3 | IP7 | 0.5408 | 0.1469 |
| IP3 | IP8 | 0.4986 | 0.1370 |
| IP4 | IP5 | 0.8744 | 0.2178 |
| IP4 | IP6 | 0.5960 | 0.1595 |
| IP4 | IP7 | 0.5295 | 0.1443 |
| IP4 | IP8 | 0.5106 | 0.1398 |
| IP5 | IP6 | 5.0815 | 0.6180 |
| IP5 | IP7 | 2.6079 | 0.4536 |
| IP5 | IP8 | 2.4152 | 0.4347 |
| IP6 | IP7 | 1.3797 | 0.3052 |
| IP6 | IP8 | 1.2859 | 0.2905 |
| IP7 | IP8 | 5.6300 | 0.6419 |

**Table S10.** Outputs of the BSSVS analysis for Cluster 7 showing the rates of transmission between sampled infected premises (IP).

| **Source** | **Target** | **Bayes Factor** | **Posterior Probability** |
|---|---|---|---|
| AIV09 Reference | IP1 | 1.2929 | 0.6743 |
| AIV09 Reference | IP2 | 0.6737 | 0.5189 |
| AIV09 Reference | IP3 | 1.0750 | 0.6325 |
| IP1 | IP2 | 8.5621 | 0.9320 |
| IP1 | IP3 | 1.9532 | 0.7577 |
| IP2 | IP3 | 0.6319 | 0.5029 |
